# A complex CTCF binding code defines TAD boundary structure and function

**DOI:** 10.1101/2021.04.15.440007

**Authors:** Li-Hsin Chang, Sourav Ghosh, Andrea Papale, Mélanie Miranda, Vincent Piras, Jéril Degrouard, Mallory Poncelet, Nathan Lecouvreur, Sébastien Bloyer, Amélie Leforestier, David Holcman, Daan Noordermeer

## Abstract

Topologically Associating Domains (TADs) compartmentalize vertebrate genomes into sub-Megabase functional neighbourhoods for gene regulation, DNA replication, recombination and repair^1-10^. TADs are formed by Cohesin-mediated loop extrusion, which compacts the DNA within the domain, followed by blocking of loop extrusion by the CTCF insulator protein at their boundaries^11-20^. CTCF blocks loop extrusion in an orientation dependent manner, with both experimental and in-silico studies assuming that a single site of static CTCF binding is sufficient to create a stable TAD boundary^21-24^. Here, we report that most TAD boundaries in mouse cells are modular entities where CTCF binding clusters within extended genomic intervals. Optimized ChIP-seq analysis reveals that this clustering of CTCF binding does not only occur among peaks but also frequently within those peaks. Using a newly developed multi-contact Nano-C assay, we confirm that individual CTCF binding sites additively contribute to TAD separation. This clustering of CTCF binding may counter against the dynamic DNA-binding kinetics of CTCF^25-27^, which urges a re-evaluation of current models for the blocking of loop extrusion^21-23^. Our work thus reveals an unanticipatedly complex code of CTCF binding at TAD boundaries that expands the regulatory potential for TAD structure and function and can help to explain how distant non-coding structural variation influences gene regulation, DNA replication, recombination and repair^5,28-34^.

---

The formation of vertebrate TADs —through continuously ongoing and energy-dependent Cohesin-mediated loop extrusion^13-17,35-37^—requires the presence of boundaries between neighbouring domains. The enrichment of CTCF binding sites at these boundaries, particularly in a convergent orientation, was identified early-on^1,3,21,23,38^. Since then, most experimental studies and *in-silico* models have assumed that a single site of correctly oriented and static CTCF binding is sufficient to create a functional TAD boundary. Indeed, the inversion of CTCF binding sites can reduce TAD insulation, albeit incompletely^21,22^. Other observations converge to challenge the static and singular nature of CTCF binding at TAD boundaries. Single-molecule imaging of CTCF binding suggests a highly dynamic process, with DNA residence time mostly in the order of seconds to minutes^25,27^. Absolute quantitation of CTCF amounts combined with mathematical modelling of residence time suggest that around half to two thirds of CTCF binding sites are occupied at any time in mouse and human cells^39,40^. These dynamic DNA binding kinetics are considerably shorter than the average association of Cohesin with chromatin, suggesting TAD boundaries may be permissive to loop extrusion ‘read through’^19,25,41^. In single-cell Hi-C and super-resolution imaging experiments, the (partial) intermingling of neighbouring TADs is commonly observed, confirming that TAD boundaries are not absolute entities^42-46^. We recently showed that in population-averaged Hi-C data most TADs are not separated by punctuated TAD boundaries but rather by extended transition zones where insulation between neighbouring domains gradually increases^19^. CTCF peaks often cluster within these transition zones near TAD boundaries^47-50^, suggesting that multiple CTCF binding sites may be required for the strong separation between neighbouring domains. Here, we used comparative genomics to analyse the functional impact of CTCF clustering in transition zones on TAD boundary structure and function in mouse embryonic stem cells (mESCs), followed by experimental validations using Nano-C, a newly developed multi-contact 3C (Chromosome Conformation Capture) approach, and dedicated polymer simulations.

## TAD boundaries are composed of clustered and strong CTCF binding peaks

To obtain an unbiased inventory of CTCF binding in mESCs, we performed CTCF ChIP-seq experiments followed by peak calling without pre-selected thresholding. To filter for *bona-fide* CTCF binding peaks, we performed CTCF motif discovery and determined the optimal significance cut-off score based on diminishing returns (Fig. 1a, elbow in the curve). Using this approach, we identified over 83,000 CTCF peaks with at least one significant CTCF binding motif (Fig. 1a, Extended Data Table 1). To determine if clustering of CTCF peaks is enriched at TAD boundaries, we intersected our list of peaks with TAD boundaries obtained after determining the insulation score of published high-resolution population-averaged Hi-C data^51,52^. This analysis confirmed that a larger fraction of CTCF peaks that clustered nearby other peaks were enriched near TAD boundaries (Fig. 1b). Consequently, over 90% of TAD boundaries contained more than one peak in the 100 kb window surrounding the Hi-C boundary (Extended Data Fig. 1a; median = 5 peaks, maximum = 24 peaks). Moreover, these peaks had a considerably higher average enrichment value as compared to peaks elsewhere in the genome (Extended Data Fig. 1b). The importance of CTCF peak clustering was confirmed by the effect on the insulation score, with the number of CTCF peaks directly scaling with insulation between neighbouring regions in the Hi-C matrix (Extended Data Fig. 1c). We conclude that the large majority of TAD boundaries are composed of multiple strong CTCF peaks within extended transition zones.

**Figure 1.**
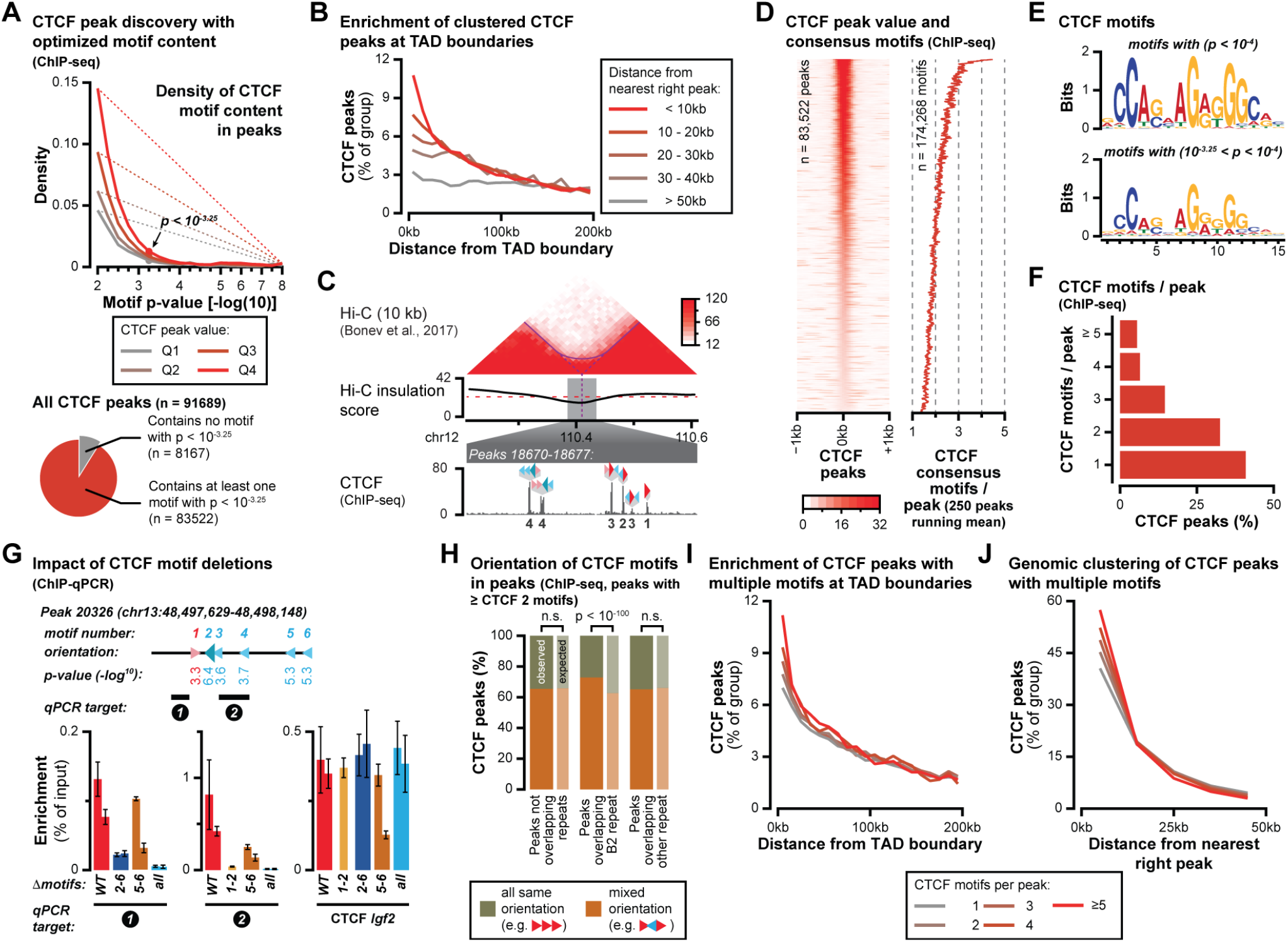
Clustering of CTCF binding within and among peaks is enriched at TAD boundaries. **A**. CTCF peak discovery based on optimal consensus-motif discovery identifies large numbers of CTCF binding peaks in mESCs. **B**. CTCF peaks that are located close to another CTCF peak are enriched closer to TAD boundaries. **C**. Example of a modular TAD boundary that appears as an extended transition zone (dotted line extending to the called TAD boundary vs. the arched line indicating extensive intermingling around the boundary. Within the transition zone, multiple CTCF peaks are present that often contain more than one CTCF consensus binding motif. Numbers below the peaks indicate the number of motifs within the selected peak. Arrows indicate the relative orientation of motifs, with the larger arrow denoting the most significant motif. **D**. Ranking of all identified CTCF peaks based on peak value (left) and the number of motifs using the same ranking (running mean). Globally, increased peak value scales with increasing numbers of CTCF motifs within peaks. **E**. CTCF binding logos for highly significant consensus motifs (top) or lower significant peaks (bottom). **F**. Distribution of CTCF consensus motifs per peak. **G**. Genome-editing experiments that remove one or several CTCF binding motifs from a peak confirm the additive contribution of individual motifs to overall CTCF enrichment as observed by ChIP qPCR. **H**. The relative orientation of CTCF motifs within peaks is mostly random, except in peaks that overlap B2 repeats. **I**. CTCF peaks that have a larger number of motifs are enriched closer to TAD boundaries. **J**. CTCF peaks that have a larger number of motifs localize closer to other CTCF peaks.

## Most CTCF peaks contain multiple consensus motifs that contribute to CTCF binding

While using our tailored identification of CTCF consensus motifs, we noticed that many CTCF peaks contained more than one significant CTCF motif (Fig. 1c, Extended Data Fig. 1d, Table 1), as previously observed^53^. Sorting both on CTCF peak enrichment value or on motif number revealed a positive correlation, suggesting that the multiple consensus motifs within the peaks contribute to peak enrichment value (Fig. 1d, Extended Data Fig. 1e). When comparing consensus motifs, we noticed that our additionally identified consensus motifs still bore the hallmarks of CTCF motifs as identified with a standard cut-off, albeit with increased promiscuity at most positions (Fig. 1e). Of the 83,000 identified CTCF peaks, nearly 60% contained more than one consensus motif (Fig. 1f; average = 2.1 motifs, maximum = 20 motifs). To confirm the contribution of multiple consensus motifs to CTCF peak value, we used CRISPR-Cas9 genome editing to remove subsets of motifs within a CTCF peak (Fig. 1g). ChIP-qPCR confirmed a partial reduction of peak enrichment when subsets of motifs were removed, as compared to the complete removal of the peaks (Fig. 1g). This result thus confirms the contribution of all motifs to overall peak enrichment. The majority of CTCF peaks are thus composed of clustered CTCF binding motifs, which adds a possibility for further regulation of CTCF binding to the DNA within these elements.

## Relative CTCF motif orientation within peaks is mostly non-biased, except in B2 family repeats

When the loop extruding Cohesin complex first encounters the N-terminus of the bound CTCF protein, it’s much more stably blocked as compared to the C-terminus, thereby explaining the orientation-dependent effect of CTCF binding^54,55^. The grouping of multiple consensus motifs within CTCF peaks may influence this process, particularly when opposing motifs are combined to create bi-directional blocking. Assessment of the relative orientation in all peaks with two or more motifs detected a significant enrichment of peaks with mixed orientations (Extended Data Fig. 2a). A more detailed analysis revealed that this enrichment is only observed for sites that overlap the B2 family of SINE-repeats, whose consensus sequence in rodent genomes contains a CTCF motif^56^. In contrast, we did not detect an enrichment for a specific relative orientation of motifs for peaks that did not cover repeats or that covered other types of repeats (Fig. 1h). Importantly though, of all the CTCF peaks in the mESC genome, about 40% contain significant binding motifs in both orientations and may thus be capable of efficient bi-directional Cohesin blocking^54,55^. Particularly in peaks that group many motifs together, the balance between motifs in either orientation can considerably deviate though (see Fig. 1c). To further determine a potential influence of B2 repeats on orientation-dependent blocking of loop extrusion, we investigated their links with CTCF binding and TAD boundaries. B2 repeats are subdivided into the B2 and B3 subtypes, and our motif discovery revealed the presence of a second CTCF motif in the consensus sequences of both subtypes. Interestingly, the relative orientation and position of the second motif is different though (Extended Data Fig. 2b). In the B2 subtypes, the second motif is positioned at around 30 bp distance in an opposite orientation relative to the first motif, whereas in the B3 subtypes the motif is in tandem and partially overlaps the first motif. Whereas we detected large numbers of significant CTCF peaks that overlapped both subtypes, their average peak values were highly different (Extended Data Fig. 2c,d). Peaks that overlapped the B2 subtypes had mostly low peak values, whereas B3 subtypes carried peaks that had values that were at least comparable to peaks that do not overlap repeats (Extended Data Fig. 2d). The mechanism underlying this difference remains to be determined, but may be linked to the more enriched AT-content in the CTCF consensus binding motifs for the B2 subtypes (Extended Data Fig. 2e). Due to relative overrepresentation of peaks overlapping the B2 repeats, these consensus motifs are identified as highly significant, yet this does not necessarily correlate with high CTCF peak value (Extended Data Fig. 2f). CTCF binding in the actively transposing B2 SINE-repeats, and particularly the tandemly orientated motifs in the strongly bound B3 subtypes, may thus provide rodent genomes with a unique evolutionary tool to add CTCF sites with either uni- or bi-directional blocking activity^57^.

## CTCF peaks with multiple motifs are enriched at TAD boundaries

To determine if clustering of motifs within peaks is a further defining characteristic of TAD boundaries, we determined the distance of CTCF peaks from TAD boundaries relative to motif number. Consistently, we found that the larger the number of CTCF-bound motifs, the larger the fraction of peaks that were located close to a TAD boundary (Fig. 1i). As a result, over 95% of TAD boundaries contained more than one CTCF-bound motif in a 100 kb window surrounding the Hi-C boundary (Extended Data Fig. 3a; median = 11 motifs, maximum = 79 motifs). Moreover, CTCF peaks with more motifs were more likely to be closer to other peaks, revealing a tendency for CTCF consensus motifs to cluster in the genome both within and between peaks (Fig. 1j). The importance of motif clustering was further confirmed by the insulation score, which scaled with motif number (Extended Data Fig. 3b,c). In summary, we conclude that TAD boundaries are of a modular nature, which is defined by a complex code of clustered CTCF consensus motifs both within and among peaks. This modular clustering of motifs may allow the blocking of Cohesin-mediated loop extrusion in both orientations at multiple sites surrounding the TAD boundary, thereby buffering against the dynamics of CTCF binding and thus improving insulation between neighbouring domains.

## Nano-C simultaneously captures 3C multi-contacts for multiple viewpoints

To test if individual CTCF sites contribute to the insulation function of modular TAD boundaries, we opted for multi-contact 3C technology, which can resolve complex patterns of chromatin organization with single-cell precision. Existing multi-contact approaches either lack the possibility for capture or can capture only a single site of interest (viewpoint)^58-62^, which limits their capacity for detailed analysis of CTCF peak clustering at selected TAD boundaries. To target multiple viewpoints within a single experiment, we therefore developed Nano-C (Fig 2a,b). Nano-C subjects a high-resolution 3C library (NlaIII digested; 206 bp average fragment length) to a newly developed method for Enrichment of Long DNA Fragments by Capture and Linear Amplification using *in-vitro* transcription (ELF-Clamp) and single-molecule direct-RNA sequencing (Fig 2a). After data analysis and stringent filtering of the reads, we obtained hundreds to thousands of multi-contacts for up to 12 viewpoints (Fig 2b-d, Extended Data Fig. 4a). Similar to other multi-contact assays that use single-molecule sequencing^59-61^, we identified considerably fewer contacts per read, as we had anticipated from the size distribution of our high-resolution 3C library (Extended Data Fig. 4b,c). We therefore visualized a 3C library generated using the NlaIII restriction enzyme by Cryo-EM (Extended Data Fig. 4d,e). We observed a mix of DNA topologies with lengths of up to 16 kb, with small fragments (< 2 kb) mostly consisting of circular and linear molecules. Unexpectedly, longer fragments consisted primarily of branched fragments, which made up 85% of the DNA content in the 3C library (Fig. 2e, Extended Data Fig. 4d,e). Proximity ligation of crosslinked DNA, at least when NlaIII digested, may thus generate unexpectedly complex topologies which will reduce the size of reads that can be generated in multi-contact 3C assays that incorporate long-read single-molecule (i.e. single-stranded) sequencing.

**Figure 2.**
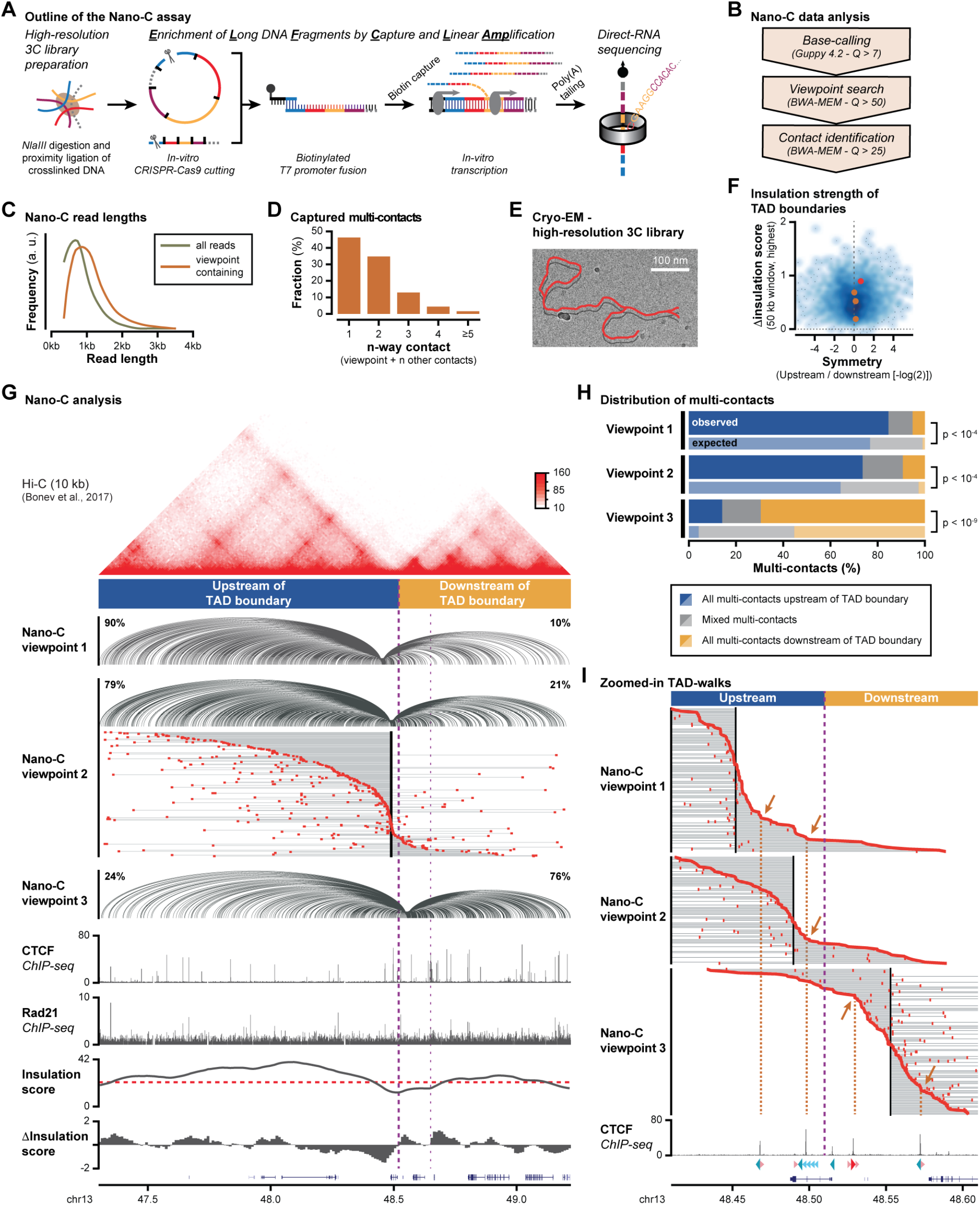
A multi-contact Nano-C assay confirms the additive contribution of CTCF peaks to TAD boundary insulation. **A**. Nano-C can enrich multiple viewpoint-containing 3C molecules using a tailored capture and linear amplification protocol (ELF-Clamp) followed by single-molecule direct-RNA sequencing to identify multi-way 3D contacts. **B**. Stringent three-step data filtering and of Nano-C reads. **C**. Nano-C read length distribution before and after data filtering. **D**. Distribution of multi-contacts captured by Nano-C. 1-way contacts represent pairwise contacts and 2-way or more contacts represent more complex 3D topologies. **E**. Example of a branched and internally looped 3C molecule, as abundantly detected by Cryo-EM. **F**. Insulation strength of all TAD boundaries in mESCs based on the derivative of the insulation score in windows 50 kb up- and downstream. Only the value for the highest window for each TAD is shown on the y-axis, whereas the symmetry is obtained by dividing the insulation strength of the upstream over the downstream boundary. Dots indicate the insulation score of the TAD boundaries analysed by Nano-C. **G**. Nano-C results for 3 viewpoints in the two TADs surrounding a strong boundary in mESCs. For all three viewpoints, spider-plots with pairwise interactions are displayed. Percentages indicate the fraction of interactions upstream or downstream of the boundary. For viewpoint 2, TAD-walks show individual multi-contact reads: each grey line represents the total span of a multi-contact read, with the viewpoint indicated as a black box and individual contacts as red boxes. Multi-contacts are sorted on the contact that is nearest to its viewpoint. The location of CTCF and Rad21 peaks (ChIP-seq) and the Hi-C insulation score (red line: cut-off) and derivative insulation score are indicated below. Previously published Hi-C data is indicated above. The thick purple line indicates the TAD boundary of interest, the smaller line indicates a second nearby boundary. **H**. Distribution of multi-contacts. Expected distributions of multi-contacts were obtained after randomizing reads up- and downstream. Significance: G-test. **I**. A zoom-in on TAD-walks in the 200 kb window surrounding the TAD boundary. Multi-contacts were sorted on the most downstream read (viewpoints located upstream of the boundary) or the most upstream read (viewpoint located downstream of the boundary). Only reads with at least one multi-contact falling within the 200 kb window are shown. Arrows and orange lines indicate cliffs that coincide with CTCF binding.

## Nano-C confirms the stepwise insulation of modular TAD boundaries

To select TAD boundaries for Nano-C in mESCs, we devised a strategy to evaluate their insulation strength based on the derivative of the insulation score in 50 kb windows up- and downstream of the boundaries, as identified by population-averaged Hi-C (Fig. 2f). This analysis revealed a wide-range of insulation strength and asymmetry in up- and downstream insulation.

We first selected a strong TAD boundary on chromosome 13 where CTCF was bound in at least four peaks and that separated a mostly transcriptionally inactive upstream TAD from a smaller intra-TAD and downstream TAD that were both transcriptionally active (Fig. 2f (red dot) and 2g, Extended Data Fig. 5). To assess the contribution of each CTCF site, we designed three Nano-C viewpoints that either flanked or were located in-between these sites. Analysis of pairwise Nano-C interactions indicated that the distribution of signal on either side of the boundary stepwise increased, with the strongest change coinciding with the TAD boundary as called from the Hi-C data (Fig. 2g; spider-plots). Similar distributions were observed with the conventional pairwise Hi-C and 4C-seq methods, confirming the robustness of the Nano-C assay (Extended Data Fig. 6). Nano-C at three weaker TAD boundaries with modular CTCF binding revealed a similar stepwise increase, albeit with less insulation between domains (Fig. 2f, brown dots, Extended Data Fig. 7a-9a).

Next, we exploited the multi-contact aspect of the Nano-C assay by filtering reads with at least two interactions within the two TADs surrounding the viewpoint. By sorting and plotting individual multi-contact reads, TAD-walks were obtained (similar to chromosome-walks^58^) that reveal the degree of insulation imposed by the selected boundary (Fig. 2g). Upon visual inspection, a consistent overrepresentation was observed for reads with both the viewpoint and all multi-contacts on the same side of the TAD boundary, which scaled with TAD boundary strength (Fig. 2g, Extended Data Fig. 7a-9a). Statistical analysis confirmed the enrichment of multi-contacts that obeyed the boundary, and revealed that the fraction of such reads was stepwise reduced depending on the location of the viewpoint relative to the boundary (Fig. 2h, Extended Data Fig. 7b-9b). Nano-C thus confirmed that TAD boundaries acted as insulating units, with the fraction of boundary-obeying multi-contacts stepwise increasing if a viewpoint was separated by a larger number of CTCF sites. Interestingly, we also found a consistent enrichment of the population of multi-contacts that are all on the other side of the boundary relative to the viewpoint (Fig. 2h, Extended Data Fig. 7b-9b). We argue that such multi-contacts may represent extruded loops that started extrusion in the TAD opposite of the viewpoint and that have partially progressed ‘read through’ of the boundary (see also below)^19,25,41^.

## Nano-C confirms the contribution of individual CTCF sites to insulation between TADs

Next, we assessed if the contribution of individual CTCF sites to TAD insulation could be directly observed in the Nano-C data. By zooming in on multi-contacts in the transition zone surrounding the TAD boundary, we observed the presence of ‘cliffs’ followed by a considerable flattening of the Nano-C interaction curve (change of angle). Such cliffs represent cases where interactions with the viewpoint locally accumulated, whereas the flattening indicates a reduced propensity for interactions. In most cases, the position of the cliffs overlapped, or were close to, CTCF sites (Fig. 2i, orange arrows, Extended Data Fig. 7c-9c). A similar reduction in contacts could be observed at the imprinted *Igf2-H19* locus, where we previously identified the presence of multiple CTCF-anchored loops to create a complex gene regulatory neighborhood^63^ (Extended Data Fig. 10). In conclusion, Nano-C allowed for a direct observation that CTCF sites reduced, but not completely blocked, contacts between the viewpoint and genomic regions on the other side of the CTCF site. Individual CTCF binding sites thus additively contributed to the overall insulating capacity of the analysed modular TAD boundaries.

## Multi-contacts spanning TAD boundaries are increased but not reorganized upon CTCF removal

To mechanistically dissect the influence of CTCF binding on Nano-C multi-contacts, we performed Nano-C experiments on mESCs that contained an Auxin-inducible CTCF degron. Previously published pairwise Hi-C studies on these cells found that insulation between TADs was greatly reduced after two days of Auxin treatment^13^. Our 2-days treatment similarly resulted in no observable remaining CTCF protein (Fig. 3a). In the absence of treatment, CTCF amounts in these cells were already strongly reduced though. To allow for an optimal comparison between the presence and absence of CTCF, we therefore limited our Nano-C comparison to WT mESCs cells.

**Figure 3.**
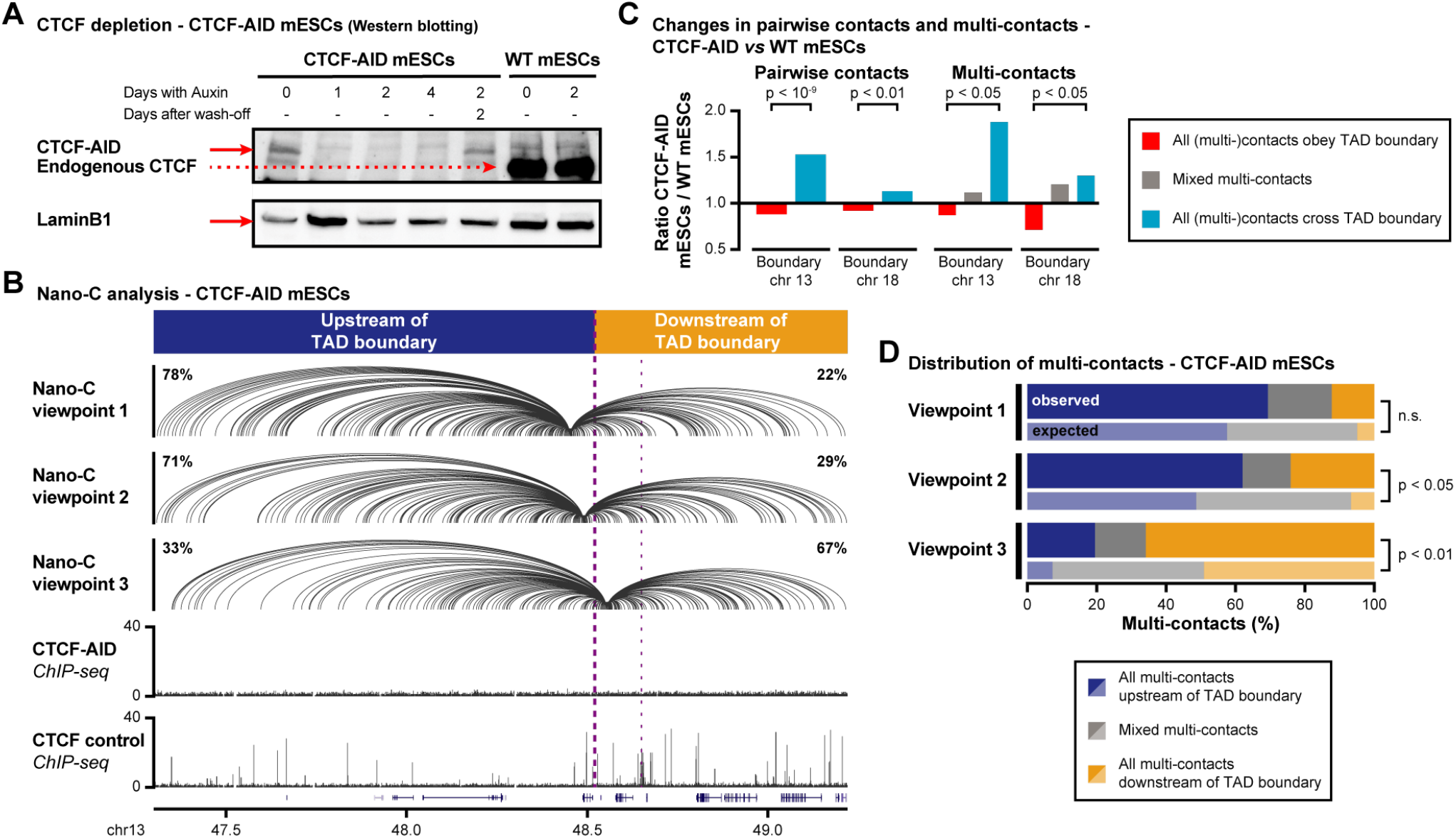
CTCF depletion perturbs the blocking of Cohesin-mediated loop extrusion at TAD boundaries. **A**. Validation of CTCF degradation upon Auxin-treatment. **B**. Nano-C results in the two TADs surrounding a strong boundary in mESCs upon CTCF depletion. Spider-plots show pairwise interactions, with percentages indicating the fraction of interactions upstream or downstream of the boundary. ChIP-seq data for CTCF binding in Auxin-treated CTCF-AID mESCs and WT mESCs is provided below. The thick purple line indicates the TAD boundary of interest, the smaller line indicates a second nearby boundary. **C**. Comparison of pairwise and multi-contact Nano-C read distribution for WT and Auxin-treated CTCF-AID mESCs at two TAD boundaries on chr. 13 and 18. Significance: G-test. **D**. Distribution of multi-contacts upon CTCF depletion. Expected distributions of multi-contacts were obtained after randomizing reads up- and downstream. Significance: G-test.

Spider-plots for pairwise Nano-C interactions at two TAD boundaries consistently confirmed that interactions that crossed the boundary had significantly increased, lending further support to the observation that CTCF is essential for correct insulation between neighbouring TADs^13,46^ (Fig. 3b,c, Extended Data Fig. 11a). Interestingly, the analysis of multi-contacts showed a more nuanced pattern. Whereas reads with both the viewpoint and all multi-contacts on the same side of the boundary were still overrepresented, their relative abundance decreased as compared to WT cells (Fig. 3c,d, Extended Data Fig. 11b). In contrast, multi-contacts that spanned the boundary were increased as compared to WT cells, which was particularly prominent for multi-contacts that all crossed the boundary relative to the viewpoint. As such, the insulating capacity of TAD boundaries was perturbed in the absence of CTCF, but this did not completely randomize 3D contacts of the viewpoints. Rather, the increase in multi-contacts that all crossed the boundary may be explained by the facilitated ‘read through’ of the boundary by the loop extrusion machinery in the absence of CTCF binding. Depletion of CTCF binding thus directly interferes with the blocking of Cohesin-mediated loop extrusion at TAD boundaries.

## A modified Randomly Cross-Linked Polymer model confirms the impact of complex TAD boundaries

To validate that the complexity of CTCF binding is an essential requisite for correct TAD structure, we used a modified Randomly Cross-Linked polymer model^64,65^ to emulate key aspects of modular TAD boundaries (Fig. 4a). Polymer simulations using a ‘naive’ model, based on random connectors that represent the Cohesin complex and that are distributed uniformly within each TAD, created a sharp boundary (Fig. 4b, left, Extended Data Fig. 12). The combined addition of three types of connectors with specific distributions were capable of reproducing the extended transition zones between TADs, while maintaining the overall insulation as observed in Hi-C maps (Fig. 4a). First, the addition of two cross-linkers that were fixed on one side next to the TAD boundary and on their other side established a contact uniformly within the same TAD, allowed to model the local concentration of cross-linker molecules (i.e. blocking of loop extrusion). Second, the addition of a gap without connectors helped to model the multiple instances of CTCF binding within the extended modular boundary. Third, the addition of moving boundaries helped to represent the dynamic nature of CTCF binding at the modular boundary.

**Figure 4.**
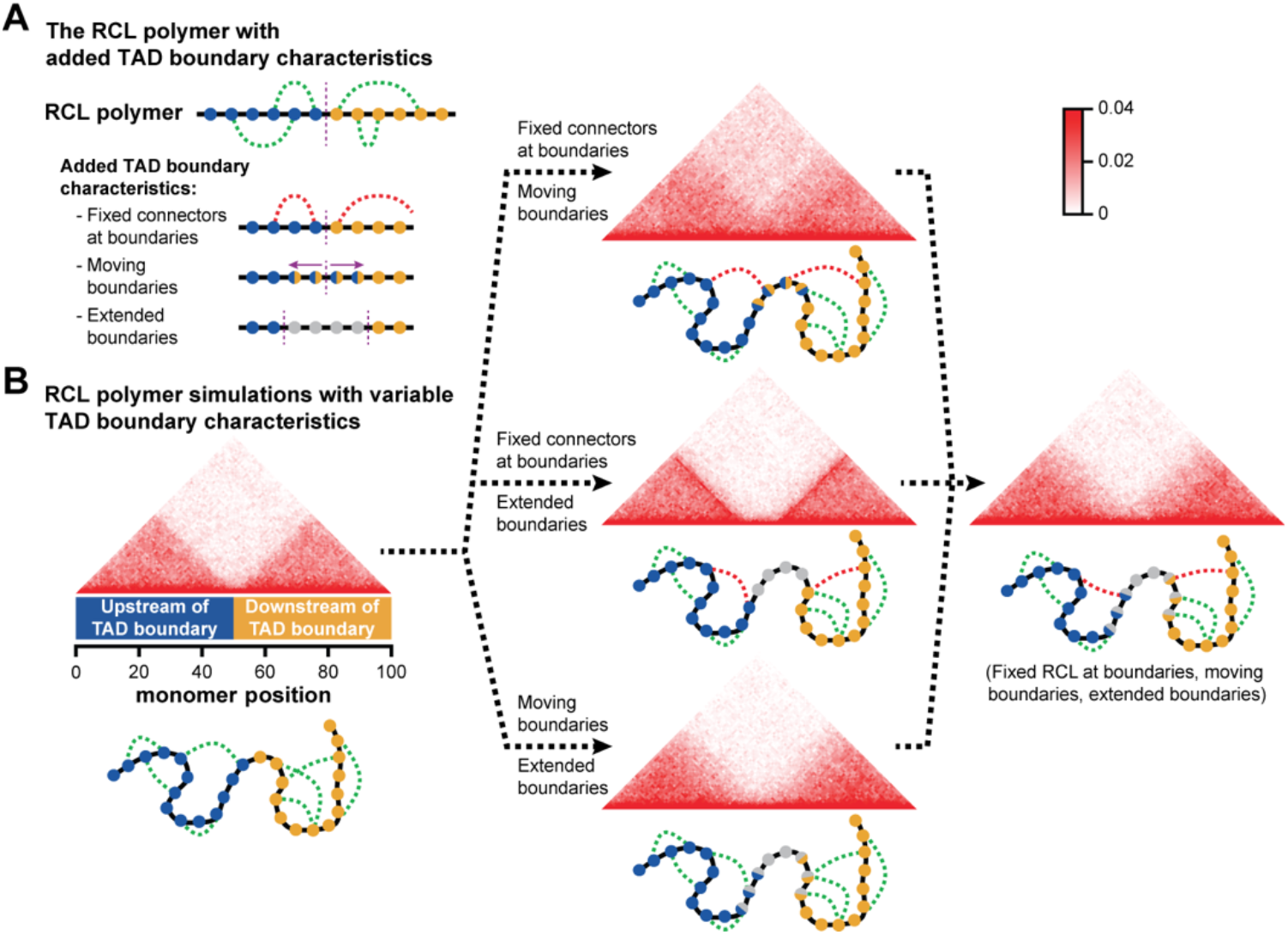
A modified Randomly Cross-Linked (RCL) polymer model replicates TAD boundary characteristics. **A**. Schematic bead-spring chain connected at random positions (green dashed lines) within the same TAD. Each sphere stands for one monomer coarse-graining a DNA fragment. Blue monomers are upstream of the TAD boundary and yellow monomers are downstream. Three additional characteristics of connectors have been added: a fixed connector to both TADs that joins the monomer next to the boundary with another random monomer within the same TAD (red dashed line), a moving boundary (two color sided monomers) and an extended boundary (grey monomers). **B**. *In-silico* Hi-C matrices obtained from polymer simulations combining the added characteristics of connectors: (left) uniformly random connectors within each TAD, (center top) fixed connectors at the boundary and a moving boundary, (center-middle) fixed connectors with an extended boundary, (center-bottom) a moving boundary with an extended boundary and (right) a combination of all three characteristics.

We found that the combination of fixed connectors at the boundary with the moving boundary resulted in a partial merger between two TADs exhibiting a fuzzy boundary (Fig. 4b, centre-top). Combination of fixed connectors with extended boundaries resulted in an overly sharp boundary alongside two dark lines corresponding to the fixed sites (Fig. 4b, centre-middle). The last pair-wise possibility, combining the moving boundary with an extended boundary, did result in an observable transition zone, but of little prominence and with increased insulation between the neighbouring TADs (Fig. 4b, centre-bottom). None of these pairwise combinations were therefore able to generate the observable transition zones while maintaining expected insulation between neighbouring TADs. Yet, the scenario where all three conditions were included recovered Hi-C maps that closely resembled the experimental ones (fig. 4b, right; e.g. relative to fig. 1c). The modelling of topological cross-linker organization, representing the Cohesin complex, including blocking at TAD boundaries both at multiple and variable positions, is thus capable of confirming the impact of dynamic and modular boundaries on TAD structure.

## Modular boundaries as regulatory units for TAD structure and function

Here, we report that most TAD boundaries are modular entities where the binding of CTCF clusters both at the level of motifs within peaks and over larger distance between peaks. Moreover, we confirm that individual CTCF peaks additively contribute to the overall insulating capacity of TAD boundaries. Compared to current models for TAD boundary function, which mostly include static and punctuated CTCF binding, this complex code of CTCF binding can provide means for fine-tuning of insulation strength (Fig. 2f), the cell-to-cell variation of TAD intermingling^45^ or TAD reorganization in a developmental context (e.g. refs. 66,67). Consequently, the spreading of TAD boundary function within extended transition zones also expands the genomic intervals where non-coding structural variation and eQTLs can modulate (sub-)TAD structuration^28,68^. This, in turn, can then create the potential for modulated or perturbed gene regulation, DNA replication, recombination and repair. Importantly though, the modular nature of TAD boundaries may create redundancies to buffer against such changes, thereby providing a potential explanation for the moderate effects on TAD structure that has been observed upon removal or inversion of individual CTCF binding sites^21,22,32-34^.

## Data availability

All unprocessed Oxford Nanopore Technologies and Illumina sequencing data (ChIP-seq, Nano-C, 4C-seq) have been deposited to the European Nucleotide Archive (EMBL-EBI ENA) under accession number PRJEB44135. Processed sequencing tracks (ChIP-seq, Nano-C, 4C-seq) have been deposited at the Mendeley Data repository: https://data.mendeley.com/datasets/g7b4z8957z/1.

## Acknowledgments

We thank Alessandra Montecucco (IGM-CNR, Pavia, Italy), Eric Joyce (University of Pennsylvania, Philadelphia, USA) and members of the Noordermeer lab for useful discussion. We thank Joke van Bemmel and Edith Heard (Institut Curie, Paris, France) for sharing the E14Tg2a.4 mESC line and Elphège P. Nora (UCSF, San Francisco, USA), Benoit Bruneau (Gladstone Institutes, San Francisco, USA) and Maxim Greenberg (Institut Jacques Monod, Paris, France) for sharing the CTCF-AID mESC line. We thank Benoit Moindrot (I2BC, Gif-sur-Yvette, France) and Marion Leleu (EPFL, Lausanne, Switzerland) for sharing the scripts for 4C-seq data analysis. This work has benefited from the facilities and expertise of the high-throughput sequencing core facility of the I2BC (Gif-sur-Yvette, France). D.H. and D.N. benefited from a collaborative PlanCancer grant (19CS145-00). This work further benefited from grants to D.N. from the Agence Nationale pour la Recherche (ANR-18-CE12-0022-02, ANR-17-CE12-0001-02, ANR-16-TERC-0027-01, ANR-14-ACHN-0009-01), the Fondation de Coopération Scientifique Campus Paris-Saclay (2015-0980i), The Fondation Bettencourt Schueller and Oxford Nanopore Technologies (MinION Early Access Programme and Direct-RNA Early Access Programme). S.G. and A.P. are supported by a postdoctoral fellowship from the Fondation pour la Recherche Médicale (Post-doctorat en France - SPF201909009328 and SPF201909009284). Cryo-EM experiments were supported by the CNRS network Microscopie Electronique et Sonde Atomique (METSA, FR CNRS 3507) to D.N. and the Investissements d’Avenir LabEx PALM (ANR-10-LABX-0039-PALM) to A. L. This project has received funding from the European Research Council (ERC) to D.H. under the European Union’s Horizon 2020 research and innovation programme (grant agreement No 882673).

## Author contributions

L.-H.C. developed Nano-C, generated Nano-C, ChIP-seq and 4C-seq data, analyzed Nano-C, ChIP-seq and 4C-seq data and wrote the manuscript.

S.G. developed the Nano-C and ChIP-seq data analysis pipeline, analyzed Nano-C, ChIP-seq and Hi-C data and wrote the manuscript.

A.P. designed the extension of the RCL polymer model, performed simulations and contributed the section on RCL modeling.

M.M. created and analyzed the mESC lines with CTCF site deletions.

V.P. developed the ChIP-seq analysis pipeline and analyzed the ChIP-seq and Hi-C data.

J.D. performed and analyzed Cryo-EM experiments.

M.P. generated ChIP-seq data.

N.L. assisted with the creation of the mESC lines with CTCF site deletions.

S.B. provided essential input for the development of Nano-C.

A.L. performed and analyzed Cryo-EM experiments.

D.H. designed the extension of the RCL polymer model and contributed the section on RCL modeling.

D.N. conceived the project, assisted in the development of Nano-C, analyzed ChIP-seq and Nano-C data, supervised L.-H.C., S.G., M.M., V.P., M.P. and N.L., and wrote the manuscript.

## Materials and methods

### ES Cell culture, CRISPR-Cas9-mediated genome editing and CTCF depletion

Feeder-independent mouse WT ESCs (male: E14Tg2a.4)^69^, their newly constructed derivaties and the CTCF-AID mESCs^13^ were cultured on gelatin-coated flasks. mESCs were grown in DMEM (GIBCO) medium supplemented with 2 mM l-Glutamine, 0.1 mM NEAA, 1 mM sodium pyruvate (GIBCO), 15% FBS (GIBCO), 10 M β-mercaptoethanol (Sigma), 1,000 U/μl of leukemia inhibitory factor (GIBCO). Medium for CTCF binding site mESCs was further supplemented with 1 mM MEK inhibitor PD0325901 (Sigma) and 3 mM GSK3 inhibitor CHIR99021 (Sigma). Cells were kept in an incubator at 37°C and 5% CO_2_.

CTCF binding sites were deleted using CRISPR-Cas9-mediated genome editing by designing gRNAs on either side of the sites (sequences in Extended Data Table 2). Sequences were cloned into the pSpCas9(BB)-2A-Puro (PX459) V2.0 plasmid (Addgene #62988; a kind gift from Feng Zhang^70^. 500,000 cells were transfected with 2.5 µg of each plasmid using Lipofectamine 3000 (Thermo Fisher Scientific). Forty eight hours after transection, cells were placed under puromycin selection (2 mg/ml) for forty eight hours. Individual colonies were seeded in 24-well plates and validated for deletions using PCR screening.

The CTCF protein was depleted in CTCF-AID mESCs by adding 500 μM of indole-3-acetic acid (IAA, a chemical analog of Auxin; Sigma-Aldrich) to the medium for 24 hrs, 48 hrs, and 96 hrs, or in the wash-off sample, 48 hrs followed by 48 hrs of medium without IAA. Depletion of CTCF was confirmed by standard Western blotting as described^13^, using 1:500 dilution of Anti-CTCF antibody (07-729, Merck-Millipore) and 1:500 dilution of Anti-Lamin B1 antibody (ab65986, Abcam).

### ChIP-Seq and ChIP-qPCR

ChIP experiments were performed as previously described^71^ with minor modifications. Cells were fixed with 2% formaldehyde solution for 10 minutes at room temperature, followed by the addition of Glycine to 0.125 M. Crosslinked chromatin was fragmented to 150-300 bp using a Covaris S220 focused-ultrasonicator device (Covaris). 10 μg of chromatin was immunoprecipitated with the one of the following antibodies: 5 μg CTCF antibody (07-729, Merck-Millipore), 5 μg RAD21 antibody (Abcam, ab992), 2 μl H3K4me3 antibody (07-473, Merck-Millipore), 5 μg H3K27ac antibody (Active Motif, 39133), 4 μg H3K36me3 antibody (Abcam, ab9050), 5 μg H3K27me3 antibody (17-622, Merck-Millipore). For quantitative comparison of CTCF binding between WT and CTCF-AID cells, 2 μg of chromatin from human MCF-7 cells was added for spike-in purposes. For ChIP-seq, indexed ChIP-seq libraries were constructed using the NEBNext Ultra II Library Prep Kit for Illumina (New England Biolabs) using the application note ‘Low input ChIP-seq’. Sequencing was done using 50 - 86 bp single-end reads on the Next-Seq 550 system (Illumina) according to the manufacturer’s instructions at the high-throughput sequencing core facility of the I2BC (Gif-sur-Yvette, France)

For ChIP-qPCR, enrichment relative to an input of immunoprecipitated chromatin fragments was determined using SsoAdvanced Universal SYBR Green Supermix (Bio-Rad) on the LightCycler480 (Roche) or the CFX384 Touch system (Bio-Rad). Primer sequences in Extended Data Table 2.

### ChIP-seq data analysis

Data was mapped to ENSEMBL Mouse assembly GRCm38 (mm10) using BWA with default parameters^72^. After removal of duplicate reads, reads with multiple alignments, and low-quality reads, densities were calculated for combined biological and technical replicates. For visualization, data densities were further binned in 200 bp windows, using the maximum density within each window.

CTCF binding peaks were identified from the combined replicates with MACS2 (version 2)^73^, using standard settings and the optimal q-value value as estimated on the basis of FDR by the tool (q < 0.05). De-novo discovery of the CTCF consensus binding motif and subsequent assignment of p-values to sequence variants was done using the MEME and FIMO tools of the MEME-suite^74,75^. We identified all binding motifs with p ≤ 10-2, allowing multiple motifs per sequence, followed by filtering on the basis of the quartiles of MACS2 peak values and the p-value of the binding motifs in steps of 0.25 -log(10) p-value. Plotting the number of all identified motifs revealed an obvious elbow in the curve at p ≤ 10^−3.25^ for all the quartiles of MACS2 peak values, where the curve bended from a mostly linear to asymptotic increase in identified motifs. CTCF binding peaks were subsequently filtered for the presence of at least one CTCF consensus binding motif with p ≤ 10^− 3.25^. A list of identified CTCF peaks and included CTCF consensus motifs is provided in Extended Data Table 1.

### Reanalysis of Hi-C data and intersection with ChIP-seq data

Hi-C sequencing data for mESCs were obtained from the GEO repository (GSE96107)^52^. Reads were mapped to ENSEMBL Mouse assembly GRCm38 (mm10) and processed to aligned reads using HiC-Pro v2.9.0 and Bowtie2 v2.3.0, with default settings to remove duplicates, assign reads to DpnII restriction fragments, and filter for valid interactions^76,77^. Hi-C interaction matrices, at 10 kb resolution, were generated from the valid interactions and were normalized using the Iterative Correction and Eigenvector decomposition method (ICE) implemented in HiC-Pro. TAD boundaries were called using TADtool^78^, with window size 500 kb and insulation score cutoff value 21.75, resulting in a high degree of genome-wide overlap with TAD borders as reported previously^52^. For the visualization of Hi-C matrices, for the creation of ‘virtual 4C plots’ and for the determination of signal densities within TADs, custom R-scripts were used.

The derivative insulation score was calculated from genome-wide insulation score at 10 kb resolution, by subtracting each bin from its downstream bin. The insulation strength at either side of individual TAD boundaries was calculated by taking the average derivative insulation score in the five bins upstream or downstream. For visualization of insulation strength the larger upstream or downstream value was retained, and for visualization of insulation asymmetry, the larger upstream or downstream value was divided by the smaller value. Coordinates of TADs, genome-wide insulation scores and genome-wide derivative insulation scores are provided in Extended Data Tables 3-5. Intersection of CTCF ChIP-seq peaks with TAD boundaries and insulation score was done using the BEDtools suite^79^.

### Nano-C

High-resolution *in-situ* Chromosome Conformation Capture (3C) libraries were generated using the conventional 3C protocol^80^, with minor modifications. 15 million cells were fixed in a 2% formaldehyde solution for 10 minutes at room temperature, followed by the addition of Glycine to 0.125 M. Cells were lysed by incubation in lysis buffer (10 mM Tris-HCl pH 8.0, 10 mM NaCl, 0.2% NP-40) supplemented with complete protease inhibitors for 20 minutes on ice. A final concentration of 0.5% SDS was added and extracts were incubated at 62°C for 10 minutes. SDS was quenched by addition of Triton X-100 to a final concentration of 1%. Chromatin was then digested with 400 U NlaIII (New England Biolabs) at 37°C for 4 hours, followed by adding another 400 U at 37°C overnight. After enzyme inactivation by incubation at 62°C for 20 minutes, DNA ligation was performed using 5 μl HC T4 DNA ligase (Promega, 50-100 U) in 1X Ligation Buffer (Promega) with 1% Triton X-100 and 10 μg/μl BSA and incubation at 16°C for 4 h. De-crosslinking was performed by adding 50 μl proteinase K (NEB) and incubation at 65°C overnight. 3C libraries were isolated by phenol-chloroform-IAA extraction followed by ethanol precipitation.

Nano-C experiments were performed using a newly developed ELF-Clamp (Enrichment of Long DNA Fragments using Capture and Linear Amplification) protocol (Fig. 2a), which consists of two selection steps for viewpoints of interest (*in-vitro* CRISPR-Cas9 cutting and site-specific fusion of a biotinylated T7 promoter) followed by specific enrichment using linear amplification (*in-vitro* transcription). The resulting RNA is subsequently characterized using direct-RNA sequencing^81^. gRNAs for *in-vitro* CRISPR-Cas9 cutting of viewpoints in the 3C libraries were produced by *in-vitro* transcription after annealing an oligo containing a T7 promoter, the specific sequence of the gRNA and a part of the common gRNA sequence, to a reverse oligo containing only the common sequence. This partially annealed template is next made fully double strand using DNA Polymerase I, Large (Klenow) Fragment (New England Biolab). *In-vitro* transcription of the resulting template to generate full-length gRNAs was performed by using the T7 RiboMAX Express Large Scale RNA Production System (Promega) followed by DNase treatment, according to the manufacturer’s instructions. gRNAs were then isolated by 1:1 phenol-chloroform-IAA extraction followed by isopropanol with sodium acetate precipitation.

CRISPR-Cas9 cutting of viewpoints in the 3C libraries was done as follows: 3300 ng each of up to 12 gRNAs was incubated with 33 pmol Alt-R S.p. Cas9 Nuclease V3 (Integrated DNA Technologies) for each gRNA in a total of 25 μl NEBuffer 3.1 (New England Biolabs) at room temperature for 10 minutes. We have noticed that in this setting, adding more than 12 gRNAs in a single experiment results in suboptimal results. 20 μg 3C library was combined with the gRNA-Cas9 complexes in a total volume of 250 μl nuclease-free water and incubated overnight at 37°C, followed by enzyme deactivation at 65°C for 30 minutes. Nuclear RNA and gRNAs were removed by adding 1250 U RNase I_f_ and incubation at 37°C for 45 minutes, followed by enzyme deactivation at 70°C for 20 minutes. The cut 3C library was purified and concentrated by adding 1 volume AMPure XP beads (Beckman Coulter) and eluted in 55 μl nuclease-free water. To repair single stranded damage, 53.5 μl of cut 3C library was mixed with 6.5 μl NEBNext FFPE Repair Buffer and 2 μl NEBNext FFPE Repair Mix (New England Biolabs), followed by incubation at 20°C for 15 minutes and addition of 3 volumes of AMPure XP beads for purification.

Site-specific fusion of a biotinylated T7 promoter was done by adding probes, on both sides of the newly generated cut site, that consisted of the following components: a first biotinylated base, a short linker sequence, the recognition site for the SfbI restriction enzyme (CCTGCAGG), the complete T7 promoter sequence (TAATACGACTCACTATAGGGAG) and a 30 bp sequence that is complementary to the sequence directly bordering the CRISPR-Cas9 cut site (see Fig. 2a). Probes were consistently designed for the both sites flanking each cut site. 0.25 μl each of the 40 mM biotinylated T7 probes on either side of each of the viewpoints were mixed and nuclease-free water was added to the final volume of 10 μl. The biotinylated probe mix was added to 28.5 μl of cut and repaired 3C library, 10 μl of 5X OneTaq buffer (New England BioLabs), 1 μl of 10 mM dNTPs and 0.5 μl of OneTaq Polymerase (New England BioLabs). To generate (partially) double stranded DNA, the reaction was incubated in a thermal cycler with the following steps: 95°C for 8 minutes, 1°C decrease per 15 sec to 65°C, 68°C for 5 minutes, rapid decrease to 4°C. Nuclease-free water was subsequently added to a final volume of 200 μl.

For each probe included in the reaction, 5 μl Dynabeads MyOne Streptavidin T1 beads (Thermo Fisher Scientific) were combined and washed according to the manufacturer’s instructions, followed by resuspension in 200 μl of B&W buffer (10 mM Tris-HCl, pH 7.0, 1mM EDTA, 2M NaCl). The total volume of beads was added to the cut and T7 promoter-fused 3C library, followed by incubation at room temperature for 30 minutes in a HulaMixer (Thermo Fisher Scientific). Beads were washed three times in 200 μl of 1X Wash buffer according to the manufacturers’ instructions, and bound DNA fragments were released by adding 43 μl nuclease-free water, 5 μl CutSmart Buffer, and 20 U SbfI (New England BioLabs), followed by incubation at 37°C for 20 minutes. The released 3C library was purified and concentrated by adding 1 volume AMPure XP beads (Beckman Coulter) and eluted in 11 μl nuclease-free water.

10 μl of the eluted 3C library was *in-vitro* transcribed using the T7 RiboMAX Express Large Scale RNA Production System (Promega) according to the manufacturer’s instructions with minor modifications: 10 μl of libraries were added to 15 μl 2x T7 RiboMax buffer, 3 μl T7 Enzyme Mix, 2 μl 5M Betaine (Sigma-Aldrich) and 1 μl SUPERase RNase Inhibitor (Thermo Fisher Scientific) and incubated at 37°C for 60 minutes. Resulting RNA was poly(A) tailed using Poly(A) Tailing Kit (Thermo Fisher Scientific) according to the manufacturer’s instructions and incubated at 37 °C for 10 minutes. The poly(A) tailed RNA was purified and concentrated by adding 100 μl of Agencourt RNAClean XP beads (Beckman Coulter,) and eluted in 12 μl nuclease-free water. RNA concentration was determined using the RNA HS Assay on a Qubit device (Thermo Fisher Scientific).

Nanopore direct-RNA sequencing libraries were prepared using the Direct-RNA Sequencing Kit, version SQK-RNA002 (Oxford Nanopore Technologies) according to the manufacturer’s instructions with minor modifications: both adaptor ligation steps were performed for 15 minutes and the reverse-transcription step was performed at 50°C for 30 minutes with 1 μl of SUPERase RNase Inhibitor supplemented. The final Agencourt RNAClean XP beads purification step was done using 24 μl of beads instead of 40 μl for a more stringent size selection. Direct-RNA sequencing was done for 48 - 72 hours, using FLO-MIN106 flowcells (R9 chemistry) on a MinION (MK 2.0) sequencing device with the MinKNOW software (Oxford Nanopore Technologies).

### Nano-C data analysis

Direct-RNA sequencing reads (fast5 format) were basecalled using Guppy 4.0.11 followed by default quality filtering (QC score > 7), which primarily removes short reads. The resulting RNA fastq files were then converted to DNA fastq using an in-house Perl script.

To reliably identify contacts within our error-prone and complex 3C reads, we devised a two-step mapping and filtering approach. Read quality was evaluated using the MinIONQC tool^82^. In the first step, reads in the DNA fastq files were mapped to a synthetic genome only consisting of the different viewpoints present in the run [viewpoint sequences obtained from ENSEMBL Mouse assembly GRCm38 (mm10)] (see Extended Data Table 6 for viewpoints contained in each run). We used BWA-MEM with default parameters^83^ on these complex 3C reads, which considerably outperformed the more commonly used Minimap2 (Nanopore Direct-RNA-seq mode). Retained reads containing the viewpoint were then filtered for high-quality mapping (MQ ≥ 50, which make up the majority of reads; see Extended Data Fig. 4a). In the second step, high-quality viewpoint-containing reads were mapped to the entire ENSEMBL Mouse assembly GRCm38 (mm10) with repeats masked (genome obtained from https://www.repeatmasker.org), using BWA-MEM with default parameters. As expected for 3C reads that are composed of fragments from different locations in the genome, we obtained both primary and supplementary mappings from within the same read. Here, we noticed a major difference between the distribution of mapping quality scores for segments that mapped to the same chromosome as the viewpoint (intra-chromosomal mappings; large majority with MQ ≥ 25) and segments that mapped to other chromosomes (inter-chromosomal mappings; large majority with MQ ≤ 25) (see Extended Data Fig. 4a). Based on the notion that the large majority of interactions in 3C experiments are intra-chromosomal^84^, we assumed that the abundant low quality inter-chromosomal mappings mostly consisted of randomly mapped reads. Individual mapping segments within our viewpoint containing reads were therefore filtered for mapping with quality scores over 25 (MQ ≥ 25).

In-house developed Perl scripts were used to assign the individual segments from each read to NlaIII fragments in the genome, to merge segments falling in the same or neighbouring NlaIII fragment and to determine the total number of multi-contacts within the reads. Reads that only mapped to the viewpoint were removed. Multi-contact information (NlaIII fragments) was organized according to the Interact Track Format, with multi-contact reads spanning over multiple lines and additional information provided in non-essential columns:

- Column 4 (Name): unique identifier assigned to each multi-contact read.
- Column 7 (Exp): multi-contacts in the read (viewpoint + n other contacts).
- Column 8 (Color): random color assigned to each multi-contact read. #000000 if the originally identified viewpoint was not identified upon mapping to the masked genome.
- Column 12 (sourceName): indication if the viewpoint was identified upon mapping to the masked genome (1: yes, 0: no).
- Column 17 (targetName): Nano-C run in which the multi-contact read was identified.

Interact files can be downloaded from the Mendeley Data repository (https://data.mendeley.com/datasets/g7b4z8957z/1) and can be visualized as Spider plots in the UCSC genome browser (http://genome.ucsc.edu). TAD-walks were generated from these Interact files after filtering for specific genomic intervals using a custom R-script.

### Determination of 3C library topology

To assess if 3C libraries generated from NlaIII-digested chromatin are composed of circular or linear molecules, we treated 200 ng of a 3C library with 0.5 U T5 exonuclease (NEB, which only degraded linear DNA) at 37°C for 60 min. 200 ng of two control plasmids (9.0 kb and 3.1kb), with and without linearization, were incubated as well. The presence or absence of degradation was analyzed by gel electrophoresis on a 0.8% agarose gel (Extended Data, figure 4c).

### Cryo-EM of vitrified 3C libraries

To determine the topology and length of the DNA molecules in a high-resolution 3C library (NlaIII used as restriction enzyme) using cryo electron microscopy, DNA molecules were trapped suspended within a thin vitreous ice layer and imaged at cryo-temperature^85^. Concentration of the 3C library was adjusted at 80 ng/μl, to ensure that DNA complexes were sufficiently concentrated for imaging but diluted enough not to overlap in most cases. 3 μl of the diluted 3C library was deposited onto a plasma-clean Quantifoil R2/2 holey carbon grid (Electron Microscopy Sciences), blotted with a filter paper for 2 seconds, and plunged into liquid ethane, using a Vitrobot Mark IV (Thermo Fisher) operated at room temperature and 100% relative humidity. Frozen grids were imaged in a JEOL 2010F transmission electron microscope equipped with a 4K Gatan Ultrascan 1000 camera at a nominal magnification of 40,000x or 50,000x. Images were recorded with a nominal defocus of 3 μm, on the camera (pixel size 0.29 or 0.236 nm) or on Kodak SO163 negative films. Negative films were developed in full strength Kodak D19 for 12 minutes, and scanned with a Coolscan 9000 (Nikon) at a resolution of 4000 pixels per inch. Images were denoised by wavelet filtration in ImageJ (‘A trous filter’ plugin, with k_1_= 20, k_n>1_= 0). DNA contour lengths projected onto the image plane (*L*_*//*_) were segmented and measured in ImageJ using the freehand line tool.

The conformation of polymer chains is modified by confinement, here in two dimensions^86,87^. The effect depends on the relative values of persistence length *l*_*p*_, contour length *L*_*c*_, and width *w* of the chain, and layer thickness *t*. Specimen thickness *t* was determined as ranging from 50 to 75 nm from stereo pairs, recorded at tilt angles ± 10°, as described^88^. As such, DNA complexes were confined within a layer whose thickness is of the order of magnitude of the persistence length of the molecule *l*_*p*_ ≈ 50 nm. As *t* ≤ 2*l*_*p*_, DNA conformation is described by the Odjik regime of confinement^87^: the relationship between the DNA length *L*_*//*_ measured on 2D projection images and the contour length *L*_*c*_ of the molecule depends on *L*_*c*_. If *L*_*c*_ ≫*t*, DNA is compressed in the z-direction and extended in xy. The projected length *L*_*//*_ relates to the contour length L_c_ as (1) *L*_*//*_ ≈ 0.9 *L*_*c*_, or, if *L*_*c*_ ≤ *t*, so that the DNA conformation is not affected by the confinement, (2) *L*_*//*_ ≈ (π/4) *L*_*c*_^89-91^. Contour length of the complexes *L*_*c*_ are calculated from the measured values of *L*_*//*_ using either (1) or (2). With a DNA chain width *w* = 2 nm ≪*t*, the projection of a linear chain on the image plane can cross itself (‘self-crossing Odijk regime’, *w* < *t* ≤ 2*l*_*p*_)^89^ and the topology of the complex cannot unambiguously be determined in all cases.

50 regions of 5 specimens frozen from the DNA library were selected for analysis, based upon the presence of well isolated complexes to exclude the possibility that neighbouring molecules overlap. Very large complexes cannot be identified without ambiguities in many cases, leading to a possible underestimation of their number. It is also possible that very small complexes (< 10 bp) present in the sample have not been detected. Complexes whose topology could not be unambiguously determined were discarded.

### 4C-seq and data analysis

Chromatin fixation, cell lysis, and 4C library preparation were done as previously described using 15 million cells per experiment^92^. NlaIII (New England Biolabs) was used as the primary restriction enzyme and DpnII (New England Biolabs) as the secondary restriction enzyme. For 4C-seq library preparation, 800 ng of 4C library was amplified using 16 individual PCR reactions with inverse primers including Illumina TruSeq adapter sequences (primers in Extended Data Table 2). Multiplexed Illumina sequencing was done using 86 bp single-end reads on the Next-Seq 550 system (Illumina) according to the manufacturer’s instructions at the high-throughput sequencing core facility of the I2BC (Gif-sur-Yvette, France).

4C-seq data were mapped to ENSEMBL Mouse assembly GRCm38 (mm10), translated to restriction fragments and smoothed (11 fragments running mean) using a stand-alone version of the 4C-seq analysis pipeline that was previously included in the HTSstation tool^93^ (scripts available upon request). For determining signal density within TADs, the raw values per restriction fragment were used. For visualization, the smoothed values were used.

For *H19* and *Igf2* 4C-seq tracks, previously published smoothed values (11 fragments running mean) were obtained from https://data.mendeley.com/datasets/6t33n6nm96/3^63^.

### Simulations using the Randomly Cross-Linked polymer model

We describe chromatin organization using our previously developed Randomly Cross-Linked (RCL) polymer model^64^. The RCL polymer has *N*_*mon*_ monomers linearly connected by harmonic springs, similar to the Rouse model^94^, and an additional *N*_*c*_ random connectors between non-sequential monomers. To represent two sequential TADs, we concatenated two blocks of RCL polymers (each of them composed by *N*_*TAD*_ monomers and *N*_*c*_ random connectors)^65,95^. To investigate which of the identified aspects of CTCF binding at TAD boundaries reproduce best the transition zones between TADs, we simulated different scenarios involving the positioning of random connectors in the RCL block-polymers.

We set *N*_*mon*_ = *2N*_*TAD*_ = 100 monomers, with *N*_*c*_ = 10 random connectors for each RCL polymer; parameters as previously estimated from Hi-C maps^65^. Subsequently, we investigated several polymer folding scenarios using numerical stochastic simulations. First, we distributed random connectors uniformly within each RCL polymer (Extended Data Fig. 12, most left panel). To simulate Cohesin blocking at a punctuated TAD boundary, we next assigned at least one connector in each TAD to join the monomer that is located directly next to the boundary with another monomer randomly chosen within the same TAD (Extended Data Fig. 12, second panel). We further expanded on this scenario with the addition of an extended boundary (i.e. the presence of a gap containing *N*_*GAP*_ monomers without random connectors) that separates the two RCL polymers (third panel). In this case we set *N*_*TAD*_ = 45 and *N*_*GAP*_ = 10. To introduce the notion of dynamic CTCF binding, and thus the potential for loop extrusion ‘read through’, we introduced a moving boundary: the size of the two RCL polymers is randomly chosen *N*_*TAD*_∈[40,60], keeping the overall size of the polymer constant. Thus, we combine the gap without random connectors (extended boundary) with a moving boundary (fourth panel). We also match the moving boundary with the presence of one random connector with one side constrained at the boundary (fifth panel). The final scenario includes all additions: the connector constrained at the boundary, the gap without random connectors and the moving boundary (sixth panel).

---

**Extended data, figure 1.**
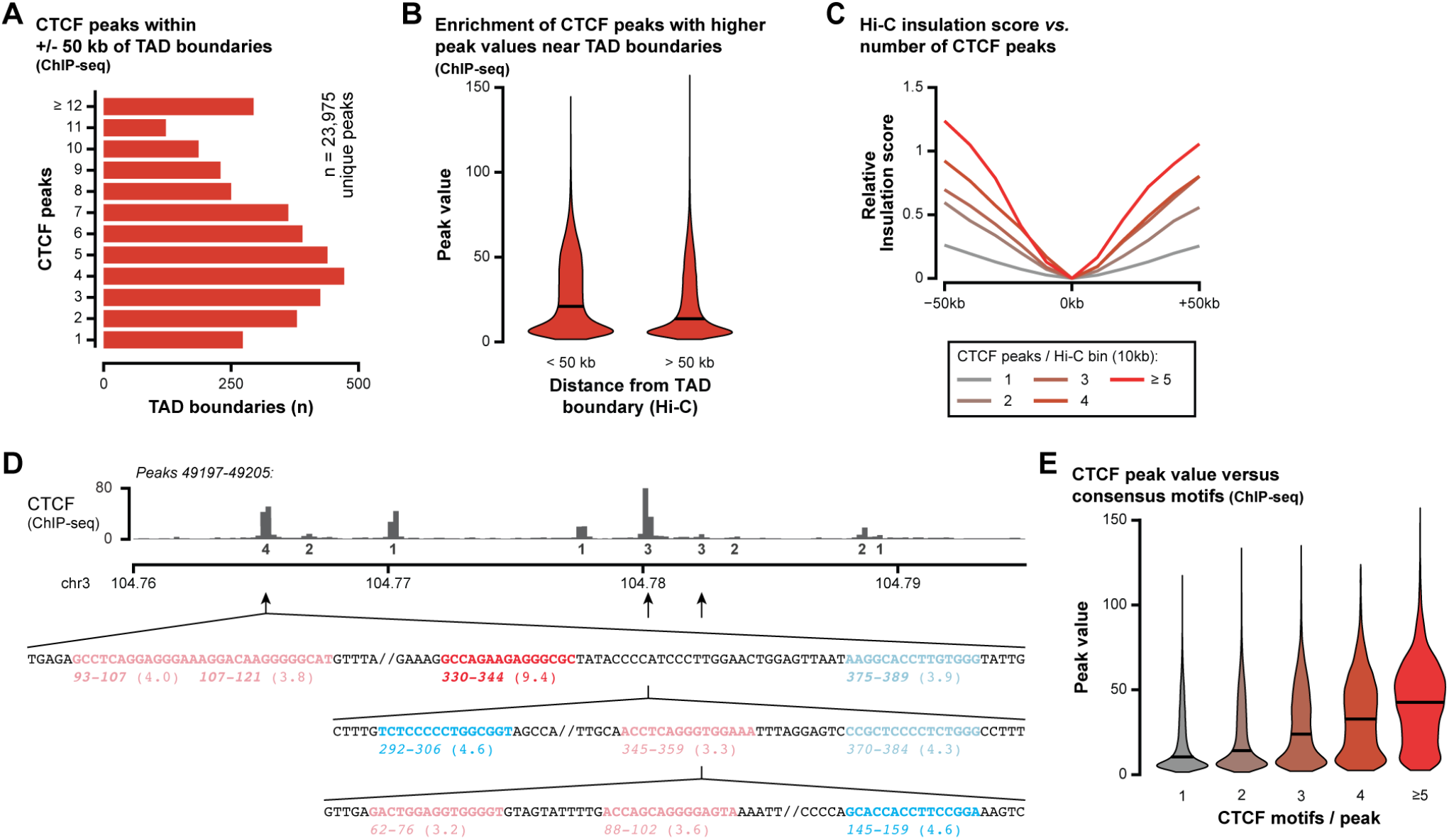
Clustering of CTCF binding near TAD boundaries and within peaks (supplemental to Figure 1) **A**. Distribution of CTCF peaks in a 100 kb window surrounding all TAD boundaries in the mESC genome, as called from population averaged Hi-C. 100 kb is the typical size of transition zones that surround TAD boundaries. **B**. Violin plots showing that peak values of CTCF peaks within 50 kb of a Hi-C boundary are, on average, increased as compared to CTCF peaks elsewhere in the genome. **C**. 10 kb Hi-C bins that carry multiple CTCF peaks, located anywhere in the mESC genome, display on average a more pronounced relative Hi-C insulation score as compared to bins that carry only one CTCF peak. **D**. Example of a cluster of CTCF binding sites, with most peaks containing multiple CTCF consensus motifs. The relative position and orientation of motifs within 3 peaks is indicated below. Red and darker blue motifs are the most significant motif within the peak. Values below each motif indicate their position within the peak and, in brackets, the significance score (-log(10) value). Red/pink motifs have a forward orientation, blue/light blue motifs have a reverse orientation. **E**. Violin plots showing that CTCF peak values are, on average, increased when more motifs are present within the peak.

**Extended data, figure 2.**
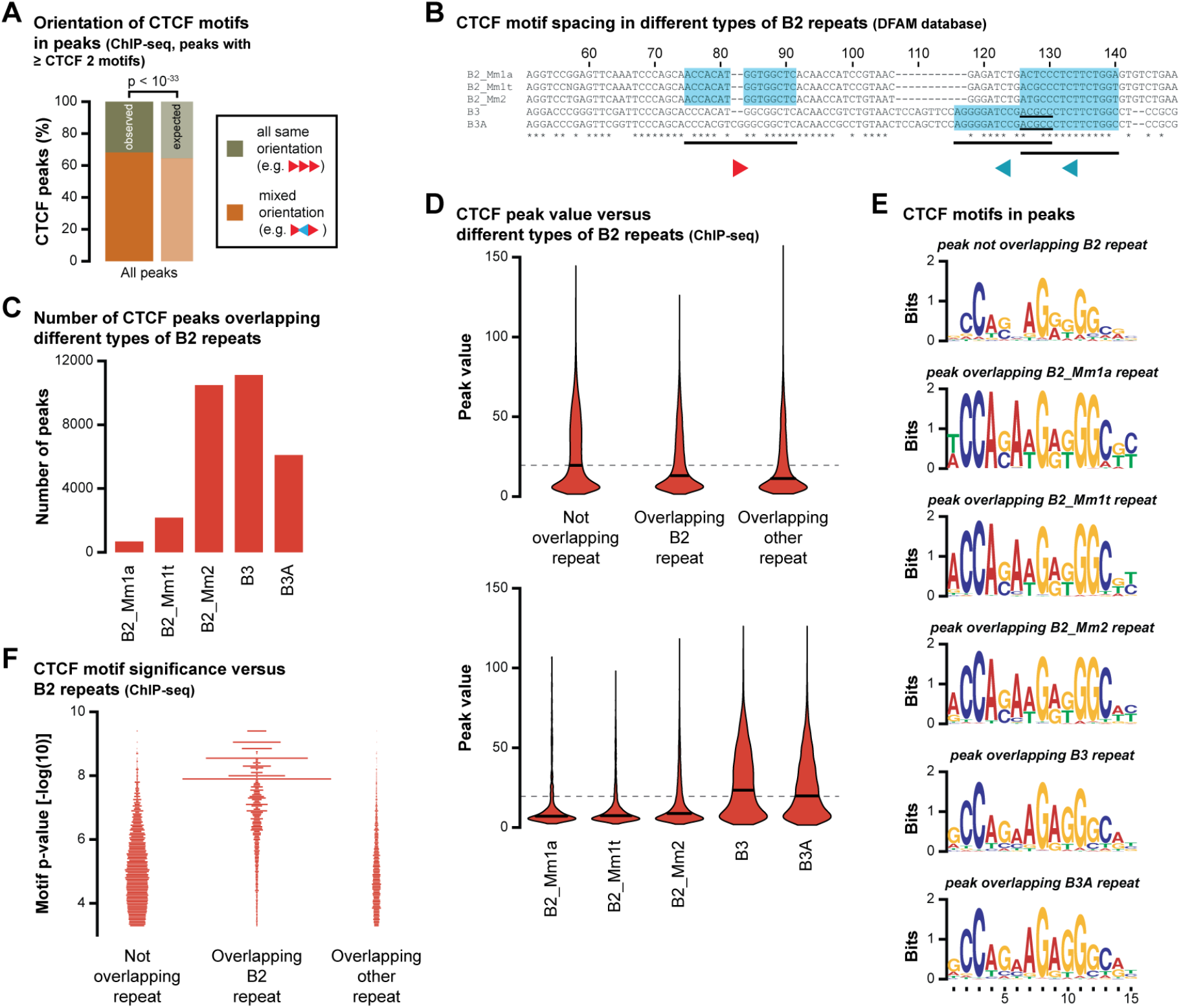
Subtypes of B2 family repeats have different CTCF binding characteristics (supplemental to Figure 1) **A**. The relative orientation of CTCF motifs within all peaks in the mESC genome is significantly enriched for peaks carrying motifs in a mixed orientation (see also Fig. 1H). **B**. The consensus sequence for B2 family repeats contain two CTCF motifs, but with a different position and orientation for different subtypes of the B2 repeat family. Blue shading highlights significant CTCF motifs. The motif that is common for all subtypes is the motif that was previously reported^56^. **C**. Total number of significant CTCF peaks in the mESC genome that overlap different subtypes of the B2 repeat family. **D**. (top) Violin plots showing that CTCF peaks that do not overlap repeats have, on average, higher peak values than peaks that overlap B2 repeats or other types of repeats. (bottom) Violin plots showing that CTCF peaks that overlap B2 subtypes have, on average, lower peak values. Peaks that overlap B3 subtypes have, on average, higher peak values that are similar to peaks that do not overlap repeats. Grey dashed line shows the median peak value of peaks that do not overlap repeats (see top panel). **E**. CTCF binding logos for CTCF motifs in peaks not overlapping B2 repeats or peaks overlapping different subtypes of B2 family repeats. **F**. Density plot showing the actual distribution of p-values for all CTCF consensus motifs, relative to repeat overlap. Due to the large overrepresentation of motifs in B2 repeats, and their relatively high degree of sequence conservation, the motifs in peaks that overlap B2 repeats are called as highly significant. Our analysis shows that this does not automatically correlate with high peak value though.

**Extended data, figure 3.**
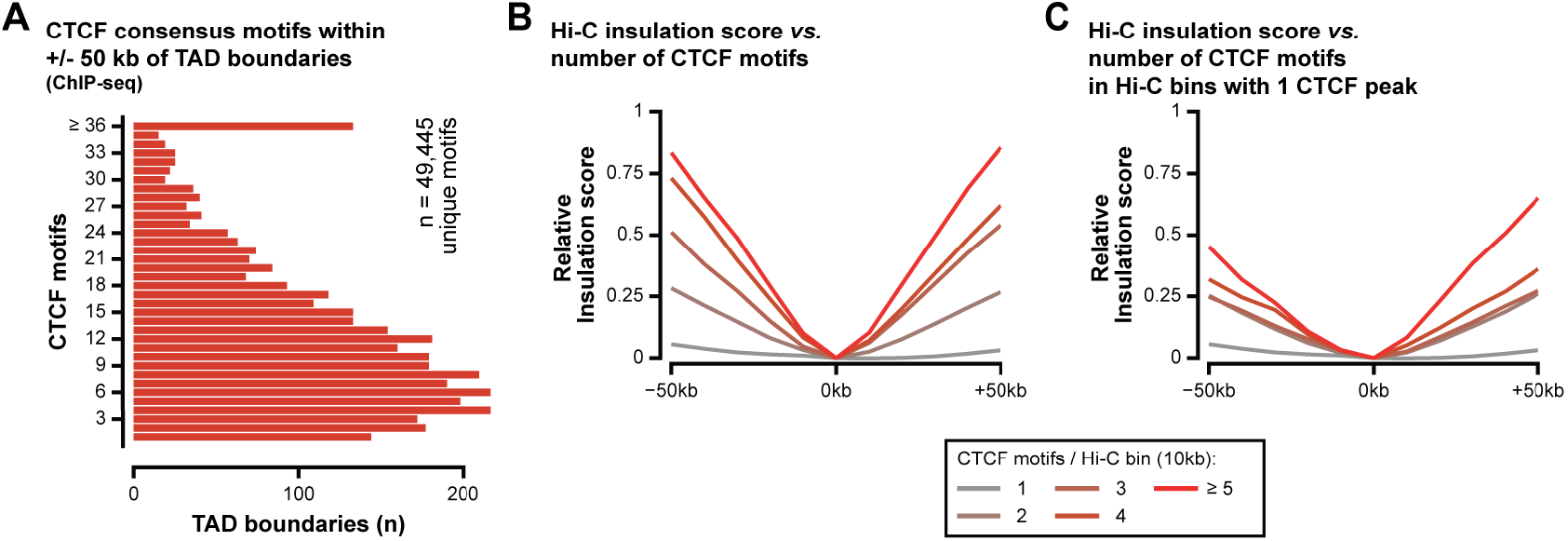
Clustering of CTCF motifs near TAD boundaries (supplemental to Figure 1) **A**. Distribution of significantly bound CTCF motifs in a 100 kb window surrounding TAD boundaries as called from population averaged Hi-C. 100 kb is the typical size of a transition zone surround TAD boundaries. **B**. 10 kb Hi-C bins that carry multiple significantly bound CTCF motifs, located anywhere in the mESC genome and in one or multiple peaks in the 10 kb bin, display on average a more pronounced relative Hi-C insulation score as compared to bins that carry only one CTCF motif. **C**. 10 kb Hi-C bins that carry a single CTCF peak with multiple significantly bound CTCF motifs, located anywhere in the mESC genome, display on average a more pronounced relative Hi-C insulation score as compared to bins that carry only one CTCF motif.

**Extended data, figure 4.**
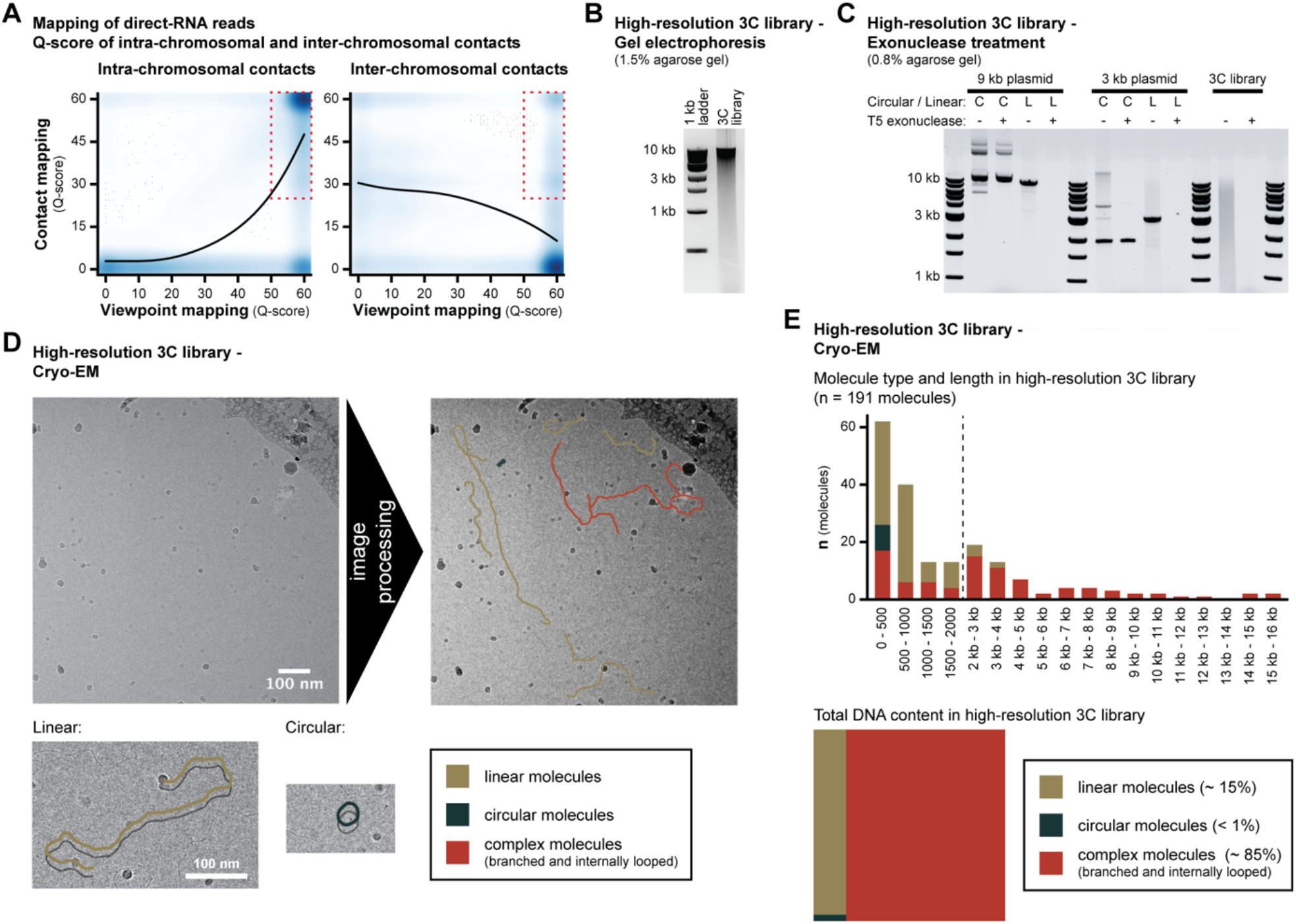
Characteristics of Nano-C reads and high-resolution 3C libraries (supplemental to Figure 2) **A**. Distribution of mapping Q-score for reads mapping both the viewpoint (second step in our data analysis; horizontal axis) and to other sites in the genome (third step in our data analysis; vertical axis). Graph on the left shows mappings where both the viewpoint and the contact mapped to the same chromosome (intra-chromosomal contact), and graph on the right shows mappings where the viewpoint and the contact mapped to different chromosomes (inter-chromosomal contact). The most prominent population of intra-chromosomal mappings had a consistently high Q-score (upper-right corner), but a smaller population was present with low Q-score for contact mapping. The most prominent population of inter-chromosomal mappings had a high quality Q-score for the viewpoint and a low Q-score for contact mapping. A smaller population was present with high Q-score for contact mapping. Considering that the mapping was done on the entire genome, we expect that specific reads should have mapped with the same Q-score anywhere in the genome. As specific contacts were enriched on the same chromosome, which represented only a small fraction of the entire genome (around 5%), we assumed that the abundance of mapping with low Q-score represented low quality non-specific mappings. For this reason, we only retained reads that were located within the red box. **B**. Gel electrophoresis of a high-resolution 3C library on a 1.5% agarose gel, showing its typical fragment distribution that appears as a relatively condensed high molecular weight band (> 10 kb). **C**. To assess if high-resolution 3C libraries generated from NlaIII-digested and proximity-ligated chromatin are of a circular or linear nature, 3C libraries were treated with T5 exonuclease, which only degrades DNA molecules that have open ends (i.e. non-circular DNA). The presence or absence of degradation was analyzed by gel electrophoresis on a 0.8% agarose gel of indicated DNA molecules, including two control plasmids with and without linearization. As for linearized plasmids, the 3C libraries also show digestion by T5 exonuclease This result confirmed that the majority of molecules in 3C libraries were not circular^62^. We noted nonetheless that the gel electrophoresis of the undigested 3C library on a 0.8% agarose gel, showed a very different distribution as on a 1.5% agarose gel (*panel B*), providing a first indication that 3C libraries were not composed of conventional linear fragments. **D**. Visualization of individual DNA molecules in a high-resolution 3C library by Cryo-EM (electron microscopy). DNA molecules were trapped suspended within a thin vitreous ice layer and imaged at cryo-temperature. Mainly, three types of topology were detected by Cryo-EM: linear molecules (highlighted in brown), circular molecules (black) and molecules with more complex topologies (branched and internally looped, red). See also Fig. 2E. **E**. Quantitation of DNA fragments and their DNA content in a high-resolution 3C library, as determined by Cryo-EM. The length of the DNA molecules was converted to base-pairs from the measured length (nm). Among the 191 molecules analyzed, the DNA molecules larger than 4 kb consisted only of complex topologies, while linear and circular molecules were mostly fragments < 2 kb (upper panel). Considering their large size, around 85% of the DNA content in the 3C library was contained within molecules with complex topologies (lower panel).

**Extended data, figure 5.**
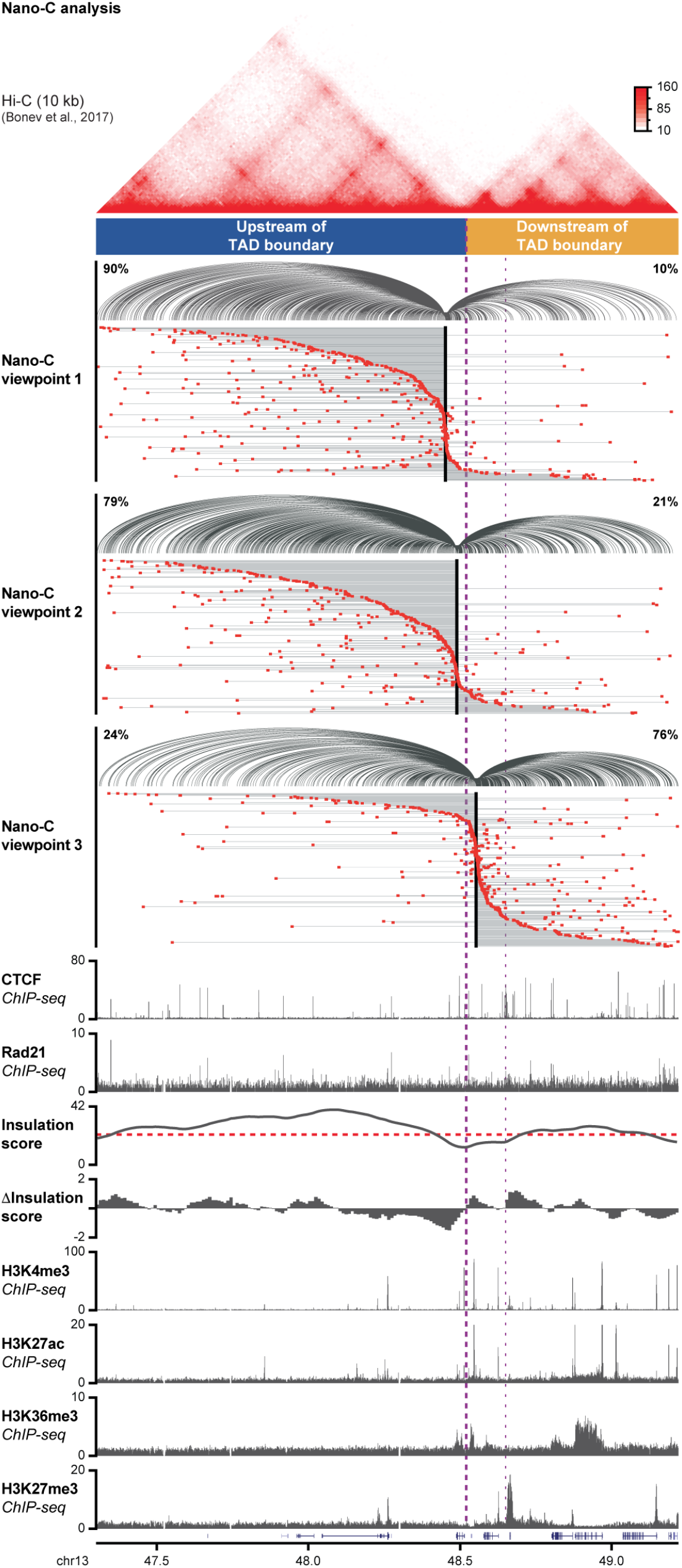
Nano-C result for three viewpoints in the 2 TADs surrounding a strong boundary on chromosome 13 in mESCs (supplemental to Figure 2) For all three viewpoints, spider-plots with pairwise interactions are displayed on top (percentages indicate the fraction of interactions upstream or downstream of the boundary) and TAD-walks are shown below. Each line in the TAD-walks represent a single multi-contact read, with the viewpoint depicted as a black box and individual contacts as red boxes. Multi-contacts are sorted on the contact that is nearest to its viewpoint. The distribution of CTCF and Rad21 peaks and H3K4me3, H3K27ac, H3K36me3, and H3K27me3 histone marks(ChIP-seq) are indicated below. The Hi-C insulation score (red line: cut-off) and derivative insulation score are indicated below. Previously published Hi-C data is indicated above^52^. The thick purple line indicates the TAD boundary of interest, the smaller line indicates a second nearby boundary. See also Fig. 2G. Notice that the TAD upstream of the boundary is mostly devoid of the H3K36me3 histone mark (indicating actively transcribed gene bodies), whereas a promoter marked by a bivalent signature (H3K4me3 and H3K27me3) is present. The TAD downstream of the boundary contains several actively transcribed genes.

**Extended data, figure 6.**
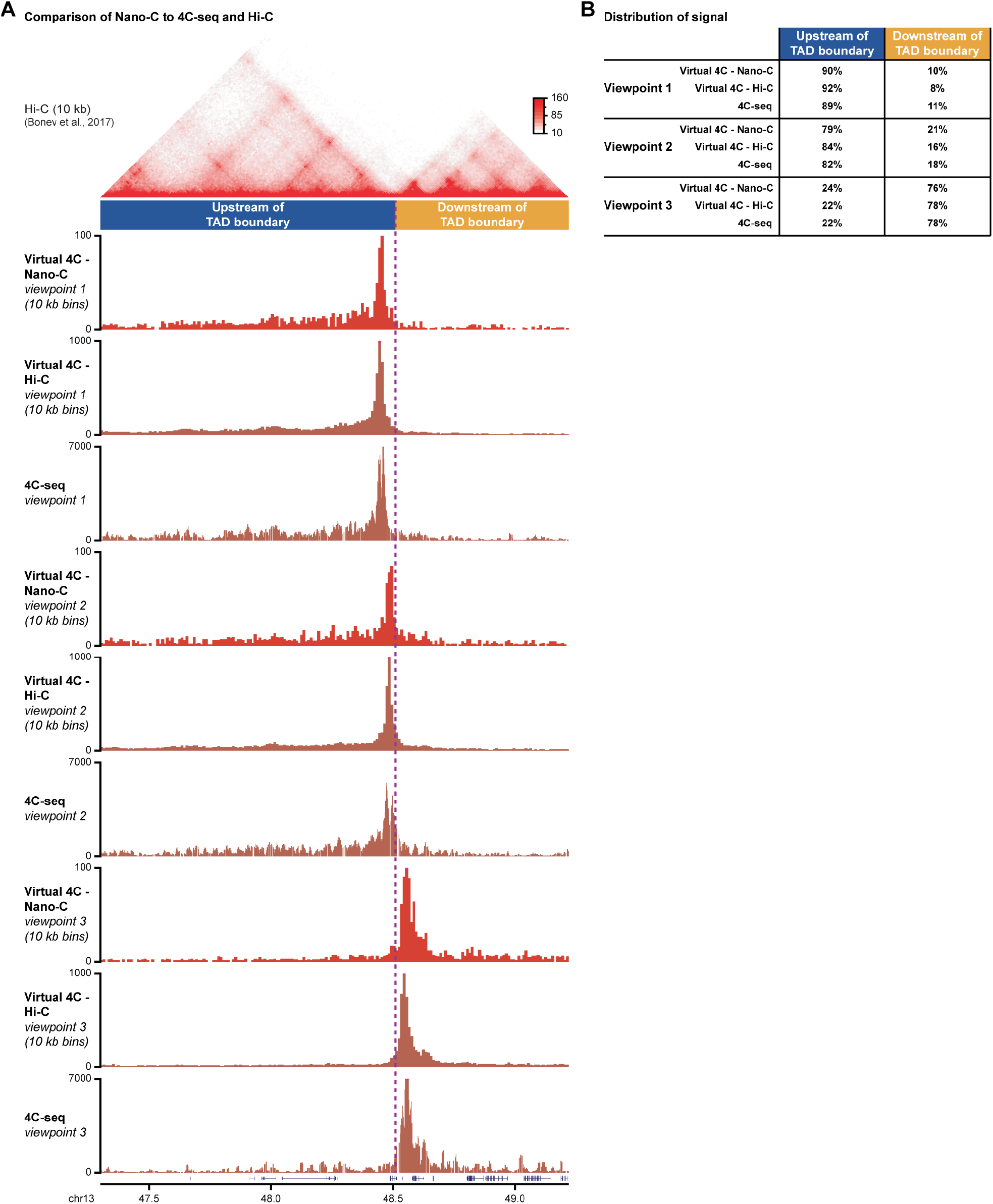
Nano-C reproduces results from pair-wise 4C-seq and Hi-C assays (supplemental to Figure 2) **A**. Comparison between Nano-C, Hi-C and 4C-seq data in the 2 TADs surrounding a strong boundary in mESCs. For the three Nano-C viewpoints on chr. 13, virtual 4C tracks^96^ were generated from the Nano-C and Hi-C data (both 10 kb resolution) and visualized with newly generated 4C-seq data (11 NlaIII fragments running-mean). The thick purple line indicates the TAD boundary of interest. Previously published Hi-C data is indicated above^52^. **B**. Distribution of signal upstream and downstream of the TAD boundary of interest for the virtual 4C and 4C-seq tracks. Distribution of signal is highly reproducible between the three assays, confirming the robustness of the Nano-C assay.

**Extended data, figure 7.**
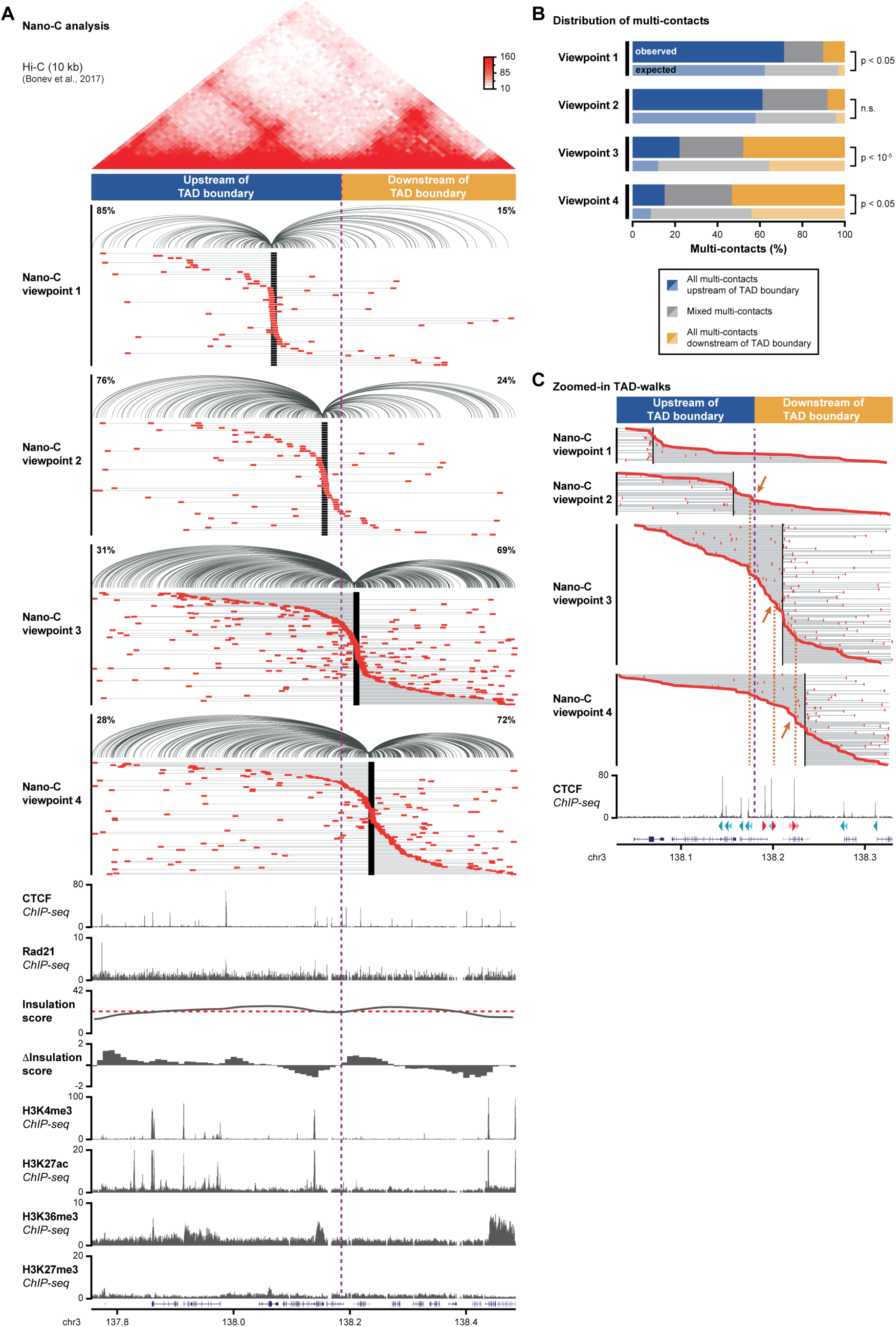
Nano-C results for four viewpoints in the 2 TADs surrounding a weaker boundary on chromosome 3 in mESCs (supplemental to Figure 2) **A**. Four viewpoints were designed around a weaker TAD boundary on chromosome 3 (see Fig. 2f, top brown dot). For all four viewpoints, spider-plots with pairwise interactions are displayed on top (percentages indicate the fraction of interactions upstream or downstream of the boundary) and TAD-walks are shown below. Multi-contacts are sorted on the contact that is nearest to its viewpoint. The distribution of CTCF and Rad21 peaks and H3K4me3, H3K27ac, H3K36me3, and H3K27me3 histone marks(ChIP-seq) are indicated below. The Hi-C insulation score (red line: cut-off) and derivative insulation score are indicated below. Previously published Hi-C data is indicated above^52^. The thick purple line indicates the TAD boundary of interest. Notice that both TADs surrounding the boundary contain several actively transcribed genes. **B**. Distribution of multi-contacts. Expected distributions of multi-contacts were obtained after randomizing reads up- and downstream. Significance: G-test. For all viewpoints, the population of mixed multi-contacts was less prevalent than expected. **C**. Zoom-in on TAD-walks in the 300 kb window surrounding the TAD boundary. Multi-contacts were sorted on the most downstream read (viewpoints located upstream of the boundary) or the most upstream read (viewpoints located downstream of the boundary). Only reads with at least one multi-contact falling within the 300 kb window are shown. Arrows and orange lines indicate cliffs that coincide with CTCF binding.

**Extended data, figure 8.**
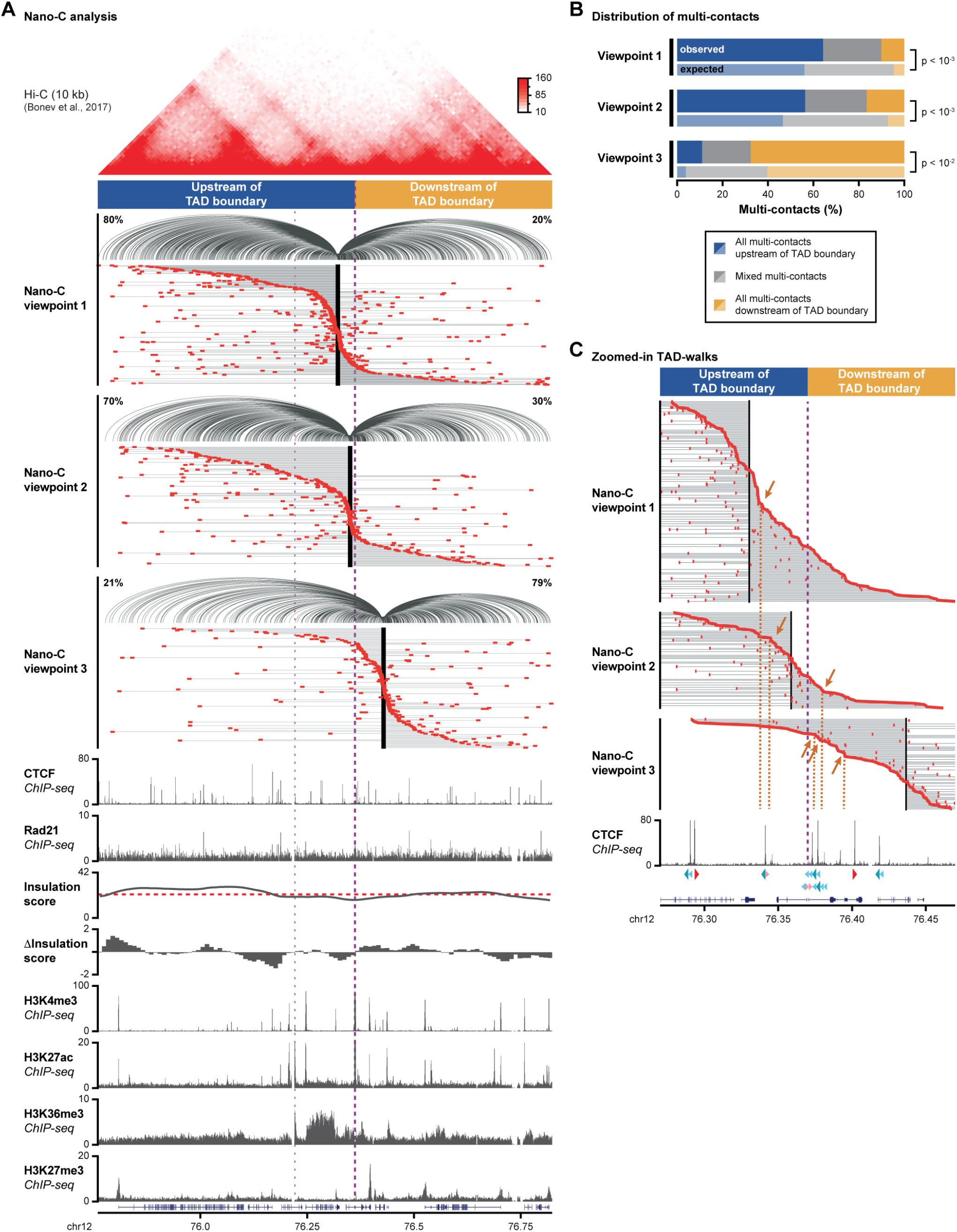
Nano-C results for three viewpoints in the 2 TADs surrounding a weaker boundary on chromosome 12 in mESCs (supplemental to Figure 2) **A**. Three viewpoints were designed around a weaker TAD boundary on chromosome 12 (see Fig. 2f, middle brown dot). For all three viewpoints, spider-plots with pairwise interactions are displayed on top (percentages indicate the fraction of interactions upstream or downstream of the boundary) and TAD-walks are shown below. Multi-contacts are sorted on the contact that is nearest to its viewpoint. The distribution of CTCF and Rad21 peaks and H3K4me3, H3K27ac, H3K36me3, and H3K27me3 histone marks(ChIP-seq) are indicated below. The Hi-C insulation score (red line: cut-off) and derivative insulation score are indicated below. Previously published Hi-C data is indicated above^52^. The thick purple line indicates the TAD boundary of interest, the smaller line indicates a second nearby boundary. Notice that the ‘inter-TAD’ upstream of the boundary contains one highly expressed gene, whereas both the TADs up- and downstream of the boundary contain inactive, bivalently marked or weakly expressed genes and promoters. **B**. Distribution of multi-contacts. Expected distributions of multi-contacts were obtained after randomizing reads up- and downstream. Significance: G-test. For all viewpoints, the population of mixed multi-contacts was less prevalent than expected. **C**. Zoom-in on TAD-walks in the 200 kb window surrounding the TAD boundary. Multi-contacts were sorted on the most downstream read (viewpoint located upstream of the boundary) or the most upstream read (viewpoints located downstream of the boundary). Only reads with at least one multi-contact falling within the 200 kb window are shown. Arrows and orange lines indicate cliffs that coincide with CTCF binding.

**Extended data, figure 9.**
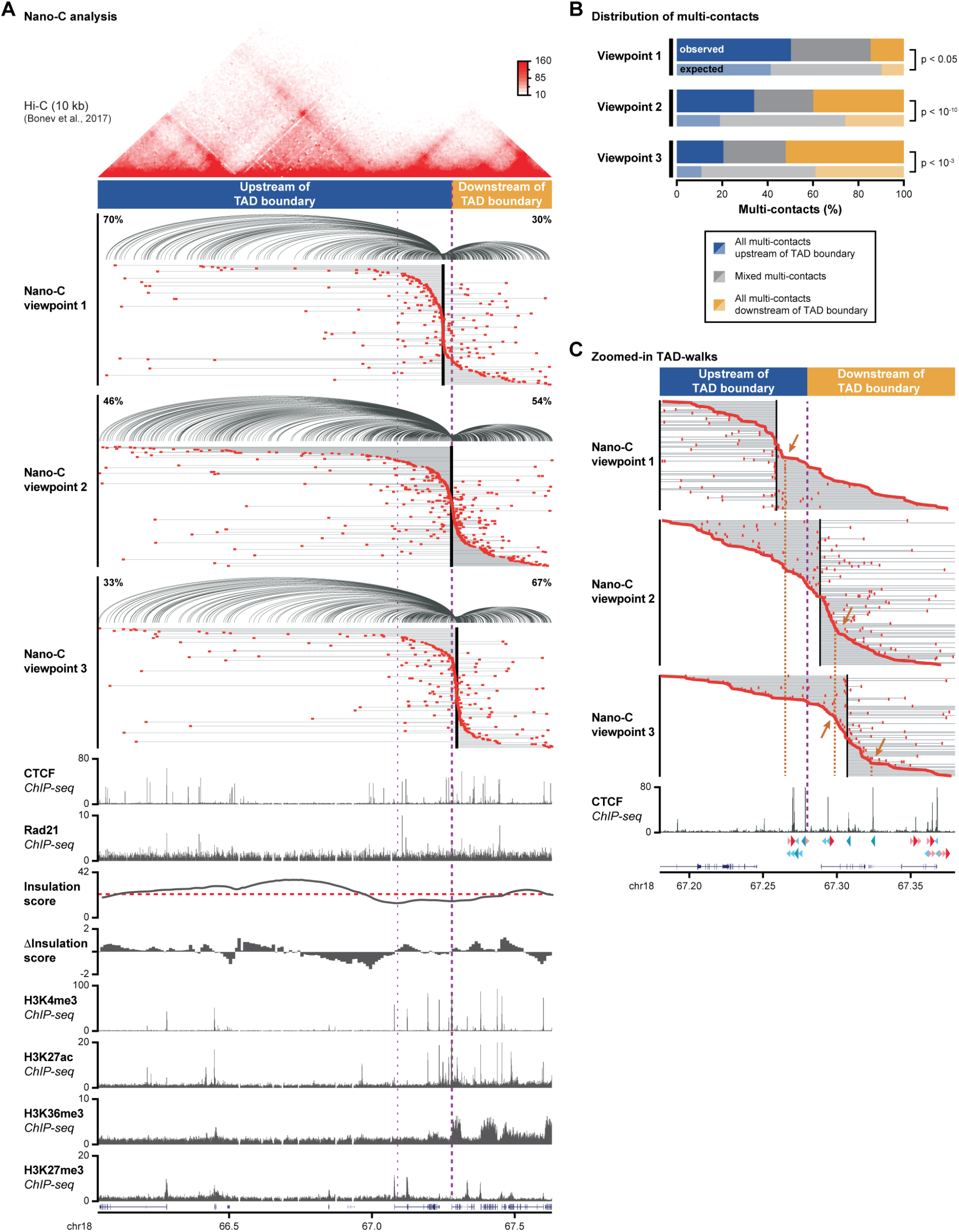
Nano-C results for three viewpoints in the 2 TADs surrounding a weak boundary on chromosome 18 in mESCs (supplemental to Figure 2) **A**. Three viewpoints were designed around a weak TAD boundary on chromosome 18 (see Fig. 2f, bottom brown dot). For all three viewpoints, spider-plots with pairwise interactions are displayed on top (percentages indicate the fraction of interactions upstream or downstream of the boundary) and TAD-walks are shown below. Multi-contacts are sorted on the contact that is nearest to its viewpoint. The distribution of CTCF and Rad21 peaks and H3K4me3, H3K27ac, H3K36me3, and H3K27me3 histone marks(ChIP-seq) are indicated below. The Hi-C insulation score (red line: cut-off) and derivative insulation score are indicated below. Previously published Hi-C data is indicated above^52^. The thick purple line indicates the TAD boundary of interest, the smaller line indicates a second nearby boundary. Notice that both the TAD and the ‘inter-TAD’ upstream of the boundary contain inactive, bivalently marked or weakly expressed genes and promoters, whereas the TAD downstream of the boundary contains several highly expressed genes. **B**. Distribution of multi-contacts. Expected distributions of multi-contacts were obtained after randomizing reads up- and downstream. Significance: G-test. For all viewpoints, the population of mixed multi-contacts was less prevalent as expected. **C**. Zoom-in on TAD-walks in the 200 kb window surrounding the TAD boundary. Multi-contacts were sorted on the most downstream read (viewpoint located upstream of the boundary) or the most upstream read (viewpoints located downstream of the boundary). Only reads with at least one multi-contact falling within the 200 kb window are shown. Arrows and orange lines indicate cliffs that coincide with CTCF binding.

**Extended data, figure 10.**
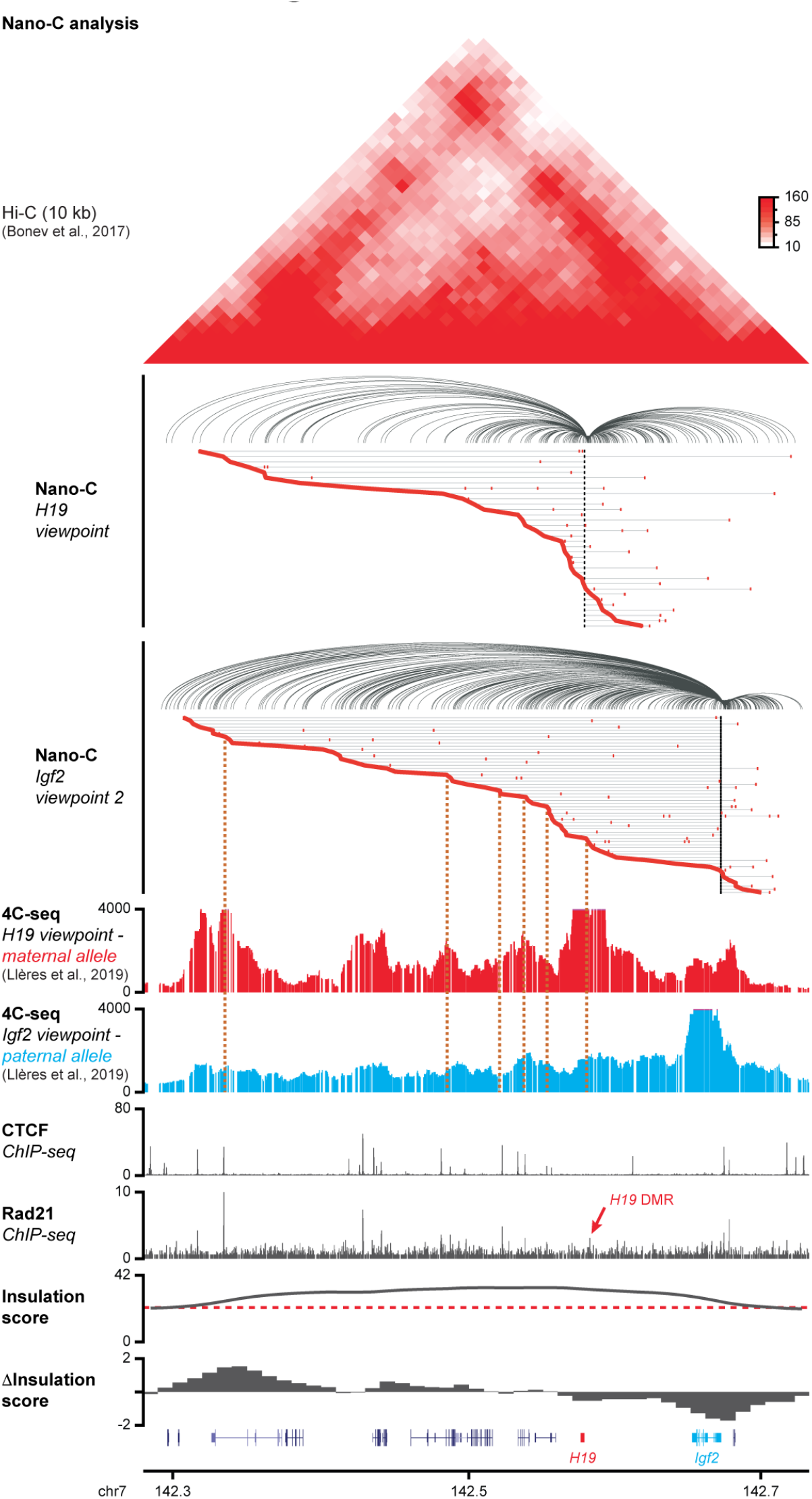
Nano-C results at the imprinted *Igf2-H19* locus in mESCs (supplemental to Figure 2) Nano-C viewpoints were designed at the *H19* DMR (differentially methylated region, which binds CTCF on the maternal allele) and at the *Igf2* promoter (which is located near to a bi-allelic CTCF site). For both viewpoints, spider-plots with pairwise interactions (top) and TAD-walks with multi-contact interactions (bottom) are displayed. Multi-contacts are sorted on the contact that is nearest to its viewpoint. Previously published allele-specific 4C-seq tracks for viewpoints at the *H19* DMR (maternal allele, with CTCF binding) and *Igf2* promoter (paternal allele, with nearby bi-allelic CTCF binding) are indicated below^63^. Non allele-specific CTCF and Rad21 peaks (ChIP-seq) are indicated below as well. Notice that CTCF binding at the *H19* DMR (red arrow), which tends to be weakly enriched^63^ was not observed, yet a relatively weak enrichment for Rad21 was visible. The Hi-C insulation score (red line: cut-off) and derivative insulation score are indicated below. Previously published Hi-C data is indicated above. Orange lines indicate cliffs that coincide with CTCF binding. Previously observed CTCF-anchored loops (4C-seq)^63^, either originating from the *H19* DMR (maternal allele) or the CTCF site downstream of the *Igf2* promoter (paternal allele) coincided with noticeable cliffs in the Nano-C data, which was particularly prominent in the *Igf2* track where we could obtain a larger number of Nano-C multi-contacts.

**Extended data, figure 11.**
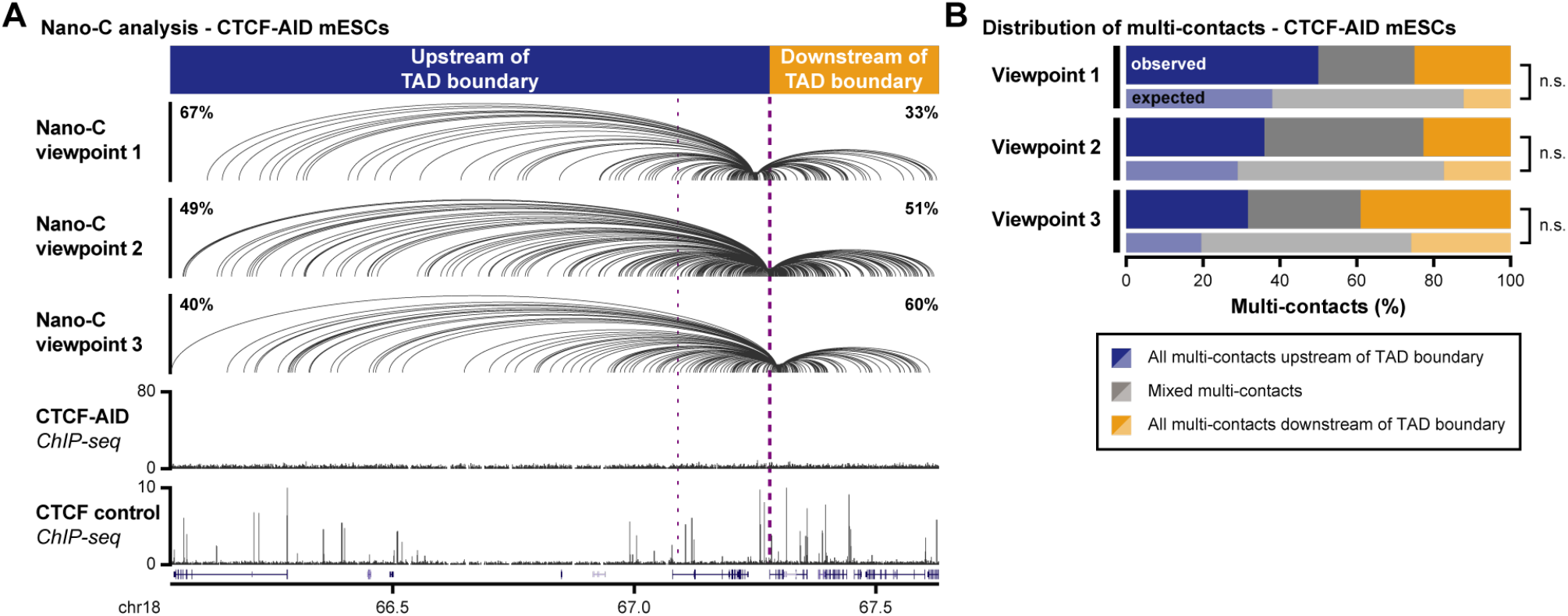
Nano-C results for three viewpoints in the 2 TADs surrounding a weak boundary on chromosome 18 in mESCs after CTCF depletion (supplemental to Figure 3) **A.**Spider-plots showing pairwise interactions for three viewpoints around a weak TAD boundary on chromosome 18, with percentages indicating the fraction of interactions upstream or downstream of the boundary (see also Extended data, figure 9). ChIP-seq data for CTCF binding in Auxin-treated CTCF-AID mESCs and WT control mESCs is provided below. The thick purple line indicates the TAD boundary of interest, the smaller line indicates a second nearby boundary. **B**. Distribution of multi-contacts. Expected distributions of multi-contacts were obtained after randomizing reads up- and downstream. Significance: G-test. Compared to WT mESCs, the population of multi-contacts for all viewpoints that obeyed the boundary was reduced (see Fig. 3C and Extended data, figure 9), but the population of mixed multi-contacts remained less prevalent than expected.

**Extended data, figure 12.**
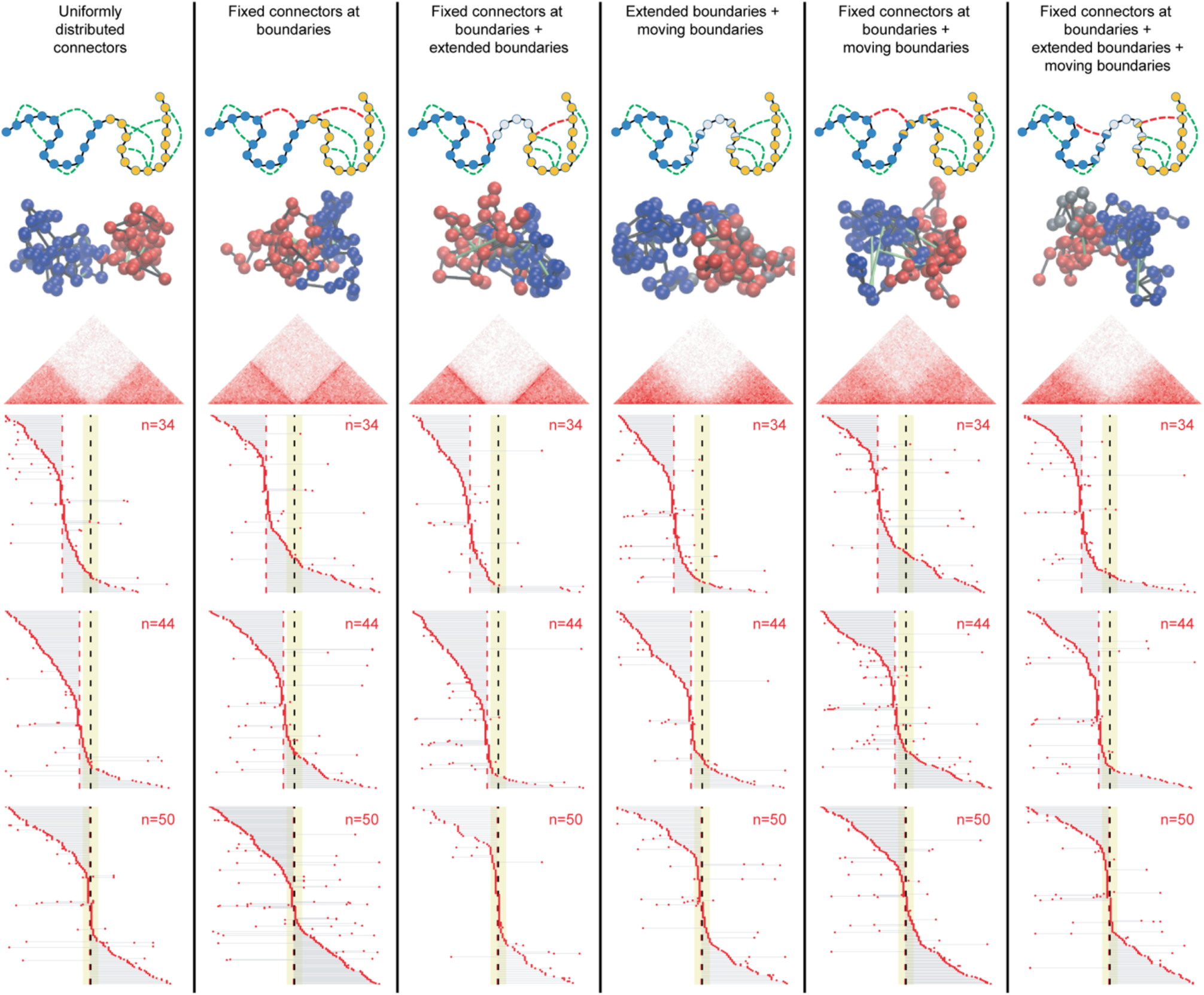
Topological configurations generated from a modified Randomly Cross-Linked (RCL) polymer model. Two sequential TADs are concatenated into two blocks of RCL polymers, each of them composed by N_TAD_ monomers linearly connected by harmonic springs with an addition of N_c_ random connectors between non-sequential monomers. Blue monomers belong to TAD 1, red/yellow monomers to TAD 2, gray monomers are the gap without random connectors. Different configurations, *in-silico* Hi-C maps and *in-silico* TAD-walks for both pairwise and multi-contacts (corresponding to monomers n=34, 44, 50, as indicated) are shown for the indicated scenarios (top). Parameters of the RCL model: b=0.18 μm, ε=0.06 μm, D=8 ·10^−3^ μm^2^/s, N_mon_=100, N_c_=10 (as determined in ref. 65). Numerical simulations are performed with time step Δt=10^−2^ s and 10^6^ integration steps. Hi-C maps and TAD walks are computed by averaging over 10^3^ polymer realizations.

## Additional files

Extended data Table 1. Identified CTCF peaks in mESCs (.xlsx)

List of identified CTCF peaks in mESCs, the CTCF consensus motifs contained within and overlap with repeats.

Extended Data Table 2. gRNA sequences, primers and probes (.pdf)

List of gRNAs used for genome editing and *in-vitro* CRISPR-Cas9 cutting (ELF-Clamp), primers used for RT-qPCR and 4C-seq, and biotinylated probes used for site-specific T7 promoter fusion (ELF-Clamp).

Extended Data Table 3. Identified TADs in mESCs (.bed)

List of start and end coordinates of TADs that were identified from reanalyzed Hi-C data^52^.

Extended Data Table 4. Insulation score in mESCs (.bedgraph)

List of insulation score for all 10kb bins in the mESC genome, as determined from reanalyzed Hi-C data^52^.

Extended Data Table 5. Derivative insulation score in mESCs (.bedgraph)

List of derivative insulation score for all 10kb bins in the mESC genome, as determined from reanalyzed Hi-C data^52^.

Extended Data Table 6. Derivative insulation score in mESCs (.xlsx)

List of NlaIII fragments used as viewpoints in each of the individual Nano-C runs.

## References

1 Dixon, J. R. et al. Topological domains in mammalian genomes identified by analysis of chromatin interactions. Nature 485, 376–380 (2012).

2 Nora, E. P., Dekker, J. & Heard, E. Segmental folding of chromosomes: a basis for structural and regulatory chromosomal neighborhoods? Bioessays 35, 818–828 (2013).

3 Dowen, J. M. et al. Control of cell identity genes occurs in insulated neighborhoods in mammalian chromosomes. Cell 159, 374–387 (2014).

4 Pope, B. D. et al. Topologically associating domains are stable units of replication-timing regulation. Nature 515, 402–405 (2014).

5 Lupianez, D. G. et al. Disruptions of topological chromatin domains cause pathogenic rewiring of gene-enhancer interactions. Cell 161, 1012–1025 (2015).

6 Ba, Z. et al. CTCF orchestrates long-range cohesin-driven V(D)J recombinational scanning. Nature 586, 305–310 (2020).

7 Collins, P. L. et al. DNA double-strand breaks induce H2Ax phosphorylation domains in a contact-dependent manner. Nat Commun 11, 3158 (2020).

8 Arnould, C. et al. Loop extrusion as a mechanism for formation of DNA damage repair foci. Nature 590, 660–665 (2021).

9 Dai, H. Q. et al. Loop extrusion mediates physiological Igh locus contraction for RAG scanning. Nature (2021).

10 Emerson, D. J. et al. Cohesin-mediated loop anchors confine the location of human replication origins. bioRxiv, 2021.2001.2005.425437 (2021).

11 Sanborn, A. L. et al. Chromatin extrusion explains key features of loop and domain formation in wild-type and engineered genomes. Proc Natl Acad Sci U S A 112, E6456–6465 (2015).

12 Fudenberg, G. et al. Formation of Chromosomal Domains by Loop Extrusion. Cell Rep 15, 2038–2049 (2016).

13 Nora, E. P. et al. Targeted Degradation of CTCF Decouples Local Insulation of Chromosome Domains from Genomic Compartmentalization. Cell 169, 930–944 e922 (2017).

14 Schwarzer, W. et al. Two independent modes of chromatin organization revealed by cohesin removal. Nature 551, 51–56 (2017).

15 Rao, S. S. P. et al. Cohesin Loss Eliminates All Loop Domains. Cell 171, 305–320 e324 (2017).

16 Wutz, G. et al. Topologically associating domains and chromatin loops depend on cohesin and are regulated by CTCF, WAPL, and PDS5 proteins. EMBO J 36, 3573–3599 (2017).

17 Vian, L. et al. The Energetics and Physiological Impact of Cohesin Extrusion. Cell 173, 1165–1178 e1120 (2018).

18 Beagan, J. A. & Phillips-Cremins, J. E. On the existence and functionality of topologically associating domains. Nat Genet 52, 8–16 (2020).

19 Chang, L. H., Ghosh, S. & Noordermeer, D. TADs and Their Borders: Free Movement or Building a Wall? J Mol Biol 432, 643–652 (2020).

20 McCord, R. P., Kaplan, N. & Giorgetti, L. Chromosome Conformation Capture and Beyond: Toward an Integrative View of Chromosome Structure and Function. Mol Cell 77, 688–708 (2020).

21 de Wit, E. et al. CTCF Binding Polarity Determines Chromatin Looping. Mol Cell 60, 676–684 (2015).

22 Guo, Y. et al. CRISPR Inversion of CTCF Sites Alters Genome Topology and Enhancer/Promoter Function. Cell 162, 900–910 (2015).

23 Vietri Rudan, M. et al. Comparative Hi-C reveals that CTCF underlies evolution of chromosomal domain architecture. Cell Rep 10, 1297–1309 (2015).

24 Mirny, L. A., Imakaev, M. & Abdennur, N. Two major mechanisms of chromosome organization. Curr Opin Cell Biol 58, 142–152 (2019).

25 Hansen, A. S., Pustova, I., Cattoglio, C., Tjian, R. & Darzacq, X. CTCF and cohesin regulate chromatin loop stability with distinct dynamics. Elife 6 (2017).

26 Gu, B. et al. Opposing Effects of Cohesin and Transcription on CTCF Organization Revealed by Super-resolution Imaging. Mol Cell 80, 699–711 e697 (2020).

27 Luan, J. et al. Distinct properties and functions of CTCF revealed by a rapidly inducible degron system. Cell Rep 34, 108783 (2021).

28 Katainen, R. et al. CTCF/cohesin-binding sites are frequently mutated in cancer. Nat Genet 47, 818–821 (2015).

29 Flavahan, W. A. et al. Insulator dysfunction and oncogene activation in IDH mutant gliomas. Nature 529, 110–114 (2016).

30 Franke, M. et al. Formation of new chromatin domains determines pathogenicity of genomic duplications. Nature 538, 265–269 (2016).

31 Hnisz, D. et al. Activation of proto-oncogenes by disruption of chromosome neighborhoods. Science 351, 1454–1458 (2016).

32 Despang, A. et al. Functional dissection of the Sox9-Kcnj2 locus identifies nonessential and instructive roles of TAD architecture. Nat Genet 51, 1263–1271 (2019).

33 Paliou, C. et al. Preformed chromatin topology assists transcriptional robustness of Shh during limb development. Proc Natl Acad Sci U S A (2019).

34 Williamson, I. et al. Developmentally regulated Shh expression is robust to TAD perturbations. Development 146, dev179523 (2019).

35 Davidson, I. F. et al. DNA loop extrusion by human cohesin. Science 366, 1338–1345 (2019).

36 Kim, Y., Shi, Z., Zhang, H., Finkelstein, I. J. & Yu, H. Human cohesin compacts DNA by loop extrusion. Science 366, 1345–1349 (2019).

37 Golfier, S., Quail, T., Kimura, H. & Brugues, J. Cohesin and condensin extrude DNA loops in a cell cycle-dependent manner. Elife 9 (2020).

38 Rao, S. S. et al. A 3D map of the human genome at kilobase resolution reveals principles of chromatin looping. Cell 159, 1665–1680 (2014).

39 Cattoglio, C. et al. Determining cellular CTCF and cohesin abundances to constrain 3D genome models. Elife 8 (2019).

40 Holzmann, J. et al. Absolute quantification of cohesin, CTCF and their regulators in human cells. Elife 8 (2019).

41 Hansen, A. S. CTCF as a boundary factor for cohesin-mediated loop extrusion: evidence for a multi-step mechanism. Nucleus 11, 132–148 (2020).

42 Nagano, T. et al. Single-cell Hi-C reveals cell-to-cell variability in chromosome structure. Nature 502, 59–64 (2013).

43 Flyamer, I. M. et al. Single-nucleus Hi-C reveals unique chromatin reorganization at oocyte-to-zygote transition. Nature 544, 110–114 (2017).

44 Bintu, B. et al. Super-resolution chromatin tracing reveals domains and cooperative interactions in single cells. Science 362 (2018).

45 Luppino, J. M. et al. Cohesin promotes stochastic domain intermingling to ensure proper regulation of boundary-proximal genes. Nat Genet 52, 840–848 (2020).

46 Szabo, Q. et al. Regulation of single-cell genome organization into TADs and chromatin nanodomains. Nat Genet 52, 1151–1157 (2020).

47 Clarkson, C. T. et al. CTCF-dependent chromatin boundaries formed by asymmetric nucleosome arrays with decreased linker length. Nucleic Acids Res 47, 11181–11196 (2019).

48 Madani Tonekaboni, S. A., Mazrooei, P., Kofia, V., Haibe-Kains, B. & Lupien, M. Identifying clusters of cis-regulatory elements underpinning TAD structures and lineage-specific regulatory networks. Genome Res 29, 1733–1743 (2019).

49 Kentepozidou, E. et al. Clustered CTCF binding is an evolutionary mechanism to maintain topologically associating domains. Genome Biol 21, 5 (2020).

50 Nanni, L., Ceri, S. & Logie, C. Spatial patterns of CTCF sites define the anatomy of TADs and their boundaries. Genome Biol 21, 197 (2020).

51 Crane, E. et al. Condensin-driven remodelling of X chromosome topology during dosage compensation. Nature 523, 240–244 (2015).

52 Bonev, B. et al. Multiscale 3D Genome Rewiring during Mouse Neural Development. Cell 171, 557–572 e524 (2017).

53 Wiehle, L. et al. DNA (de)methylation in embryonic stem cells controls CTCF-dependent chromatin boundaries. Genome Res 29, 750–761 (2019).

54 Li, Y. et al. The structural basis for cohesin-CTCF-anchored loops. Nature 578, 472–476 (2020).

55 Nora, E. P. et al. Molecular basis of CTCF binding polarity in genome folding. Nat Commun 11, 5612 (2020).

56 Schmidt, D. et al. Waves of retrotransposon expansion remodel genome organization and CTCF binding in multiple mammalian lineages. Cell 148, 335–348 (2012).

57 Gagnier, L., Belancio, V. P. & Mager, D. L. Mouse germ line mutations due to retrotransposon insertions. Mob DNA 10, 15 (2019).

58 Olivares-Chauvet, P. et al. Capturing pairwise and multi-way chromosomal conformations using chromosomal walks. Nature 540, 296–300 (2016).

59 Allahyar, A. et al. Enhancer hubs and loop collisions identified from single-allele topologies. Nat Genet 50, 1151–1160 (2018).

60 Oudelaar, A. M. et al. Single-allele chromatin interactions identify regulatory hubs in dynamic compartmentalized domains. Nat Genet 50, 1744–1751 (2018).

61 Ulahannan, N. et al. Nanopore sequencing of DNA concatemers reveals higher-order features of chromatin structure. bioRxiv, 833590 (2019).

62 Tavares-Cadete, F., Norouzi, D., Dekker, B., Liu, Y. & Dekker, J. Multi-contact 3C reveals that the human genome during interphase is largely not entangled. Nat Struct Mol Biol 27, 1105–1114 (2020).

63 Lleres, D. et al. CTCF modulates allele-specific sub-TAD organization and imprinted gene activity at the mouse Dlk1-Dio3 and Igf2-H19 domains. Genome Biology 20, 272 (2019).

64 Shukron, O. & Holcman, D. Statistics of randomly cross-linked polymer models to interpret chromatin conformation capture data. Physical Review E 96, 012503 (2017).

65 Shukron, O. & Holcman, D. Transient chromatin properties revealed by polymer models and stochastic simulations constructed from Chromosomal Capture data. PLoS computational biology 13, e1005469 (2017).

66 Andrey, G. et al. A switch between topological domains underlies HoxD genes collinearity in mouse limbs. Science 340, 1195 (2013).

67 Rodriguez-Carballo, E. et al. The HoxD cluster is a dynamic and resilient TAD boundary controlling the segregation of antagonistic regulatory landscapes. Genes Dev 31, 2264–2281 (2017).

68 Delaneau, O. et al. Chromatin three-dimensional interactions mediate genetic effects on gene expression. Science 364 (2019).

## Additional references

69 Doetschman, T. et al. Targetted correction of a mutant HPRT gene in mouse embryonic stem cells. Nature 330, 576–578 (1987).

70 Ran, F. A. et al. Genome engineering using the CRISPR-Cas9 system. Nat Protoc 8, 2281–2308 (2013).

71 Noordermeer, D. et al. Temporal dynamics and developmental memory of 3D chromatin architecture at Hox gene loci. Elife 3, e02557 (2014).

72 Li, H. & Durbin, R. Fast and accurate short read alignment with Burrows-Wheeler transform. Bioinformatics (Oxford, England) 25, 1754–1760 (2009).

73 Zhang, Y. et al. Model-based analysis of ChIP-Seq (MACS). Genome Biol 9, R137 (2008).

74 Bailey, T. L. et al. MEME SUITE: tools for motif discovery and searching. Nucleic Acids Res 37, W202–208 (2009).

75 Grant, C. E., Bailey, T. L. & Noble, W. S. FIMO: scanning for occurrences of a given motif. Bioinformatics (Oxford, England) 27, 1017–1018 (2011).

76 Langmead, B. & Salzberg, S. L. Fast gapped-read alignment with Bowtie 2. Nat Methods 9, 357–359 (2012).

77 Servant, N. et al. HiC-Pro: an optimized and flexible pipeline for Hi-C data processing. Genome Biol 16, 259 (2015).

78 Kruse, K., Hug, C. B., Hernandez-Rodriguez, B. & Vaquerizas, J. M. TADtool: visual parameter identification for TAD-calling algorithms. Bioinformatics (Oxford, England) 32, 3190–3192 (2016).

79 Quinlan, A. R. & Hall, I. M. BEDTools: a flexible suite of utilities for comparing genomic features. Bioinformatics (Oxford, England) 26, 841–842 (2010).

80 Splinter, E., Grosveld, F. & de Laat, W. 3C technology: analyzing the spatial organization of genomic loci in vivo. Methods Enzymol 375, 493–507 (2004).

81 Garalde, D. R. et al. Highly parallel direct RNA sequencing on an array of nanopores. Nat Methods 15, 201–206 (2018).

82 Lanfear, R., Schalamun, M., Kainer, D., Wang, W. & Schwessinger, B. MinIONQC: fast and simple quality control for MinION sequencing data. Bioinformatics (Oxford, England) 35, 523–525 (2019).

83 Li, H. Aligning sequence reads, clone sequences and assembly contigs with BWA-MEM. arXiv (2013). <https://arxiv.org/abs/1303.3997>.

84 Lieberman-Aiden, E. et al. Comprehensive mapping of long-range interactions reveals folding principles of the human genome. Science 326, 289–293 (2009).

85 Adrian, M. et al. Direct visualization of supercoiled DNA molecules in solution. EMBO J 9, 4551–4554 (1990).

86 Daoud, M. & De Gennes, P. G. Statistics of macromolecular solutions trapped in small pores. J. Phys. France 38, 85–93 (1977).

87 Odijk, T. The statistics and dynamics of confined or entangled stiff polymers. Macromolecules 16, 1340–1344 (1983).

88 Marko, M. & Hsieh, C. E. Three-dimensional cryotransmission electron microscopy of cells and organelles. Methods Mol Biol 369, 407–429 (2007).

89 Dai, L., Jones, J. J., van der Maarel, J. R. C. & Doyle, P. S. A systematic study of DNA conformation in slitlike confinement. Soft Matter 8, 2972–2982 (2012).

90 Micheletti, C. & Orlandini, E. Numerical Study of Linear and Circular Model DNA Chains Confined in a Slit: Metric and Topological Properties. Macromolecules 45, 2113–2121 (2012).

91 Tree, D. R., Reinhart, W.y.F. & Dorfman, K. D. The Odijk Regime in Slits. Macromolecules 47, 3672–3684 (2014).

92 Matelot, M. & Noordermeer, D. Determination of High-Resolution 3D Chromatin Organization Using Circular Chromosome Conformation Capture (4C-seq). Methods Mol Biol 1480, 223–241 (2016).

93 David, F. P. et al. HTSstation: a web application and open-access libraries for high-throughput sequencing data analysis. PLoS One 9, e85879 (2014).

94 Doi, M. & Edwards, S. F.The Theory of Polymer Dynamics. Vol. 73 (Clarendon Press, 1988).

95 Shukron, O., Piras, V., Noordermeer, D. & Holcman, D. Statistics of chromatin organization during cell differentiation revealed by heterogeneous cross-linked polymers. Nat Commun 10, 2626 (2019).

96 Sexton, T. et al. Three-dimensional folding and functional organization principles of the Drosophila genome. Cell 148, 458–472 (2012).

