## Extended Data Table 2 for "A complex CTCF binding code defines TAD boundary structure and function"

### gRNA sequences, primers and probes

#### CRISPR-Cas9 gRNAs

***Genome editing:***

| **CTCF motif deletion** | **Sequences (5’-3’)** | |
|---|---|---|
| Δ all motifs (1-6) | Upstream gRNA: | TAACACTACCTAATTGCCAG |
| | Downstream gRNA: | GCTCTTCTGTAGGCCTTCCG |
| Δ motifs 1-2 | Upstream gRNA: | TAACACTACCTAATTGCCAG |
| | Downstream gRNA: | GGATGAACAAGCCTGCCCCC |
| Δ motifs 2-6 | Upstream gRNA: | GGATGAACAAGCCTGCCCCC |
| | Downstream gRNA: | GCTCTTCTGTAGGCCTTCCG |
| Δ motifs 5-6 | Upstream gRNA: | GCCACCCTTCTAGACCGACT |
| | Downstream gRNA: | GCTCTTCTGTAGGCCTTCCG |

***ELF-Clamp: gRNAs for in vitro cutting:***
*Outline of the template for in-vitro transcription:*

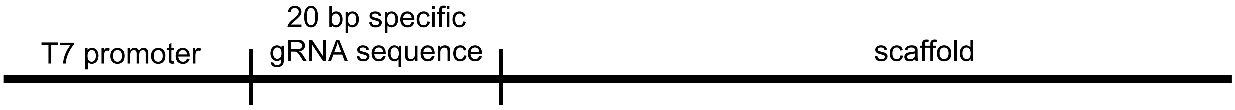


| **Primer design for *in-vitro* gRNA production** | **Sequences (5’-3’)** |
|---|---|
| T7 promoter forward primer | TAATACGACTCACTATAGGGAG - *20bp specific gRNA sequence* - GTTTTAGAGCTAGAAATA |
| Scaffold universal reverse primer | AAAAAAAGCACCGACTCGGTGCCACTTTTTCAAGTTGATAACGGA CTAGCCTTATTTTAACTTGCTATTTCTAGCTCTAAAAC |

| **Viewpoint** | **Sequences (5’-3’)** |
|---|---|
| chr13_VP1 | GAAAGGCTGCTGCTGACGTT |
| chr13_VP2 | TGAACTTCTGCCTGAAGCGC |
| chr13_VP3 | CTTATGGACACCCAGCGCCG |
| chr3_VP1 | AGAGTTGCTTGGCAGACCTG |
| chr3_VP2 | AGGGGCCGTTCCTGCTCACA |
| chr3_VP3 | ATGCTAGCAGAAGTGTCTGT |
| chr3_VP4 | CCATCGTGTGAGGCTTTGGC |
| chr12_VP1 | CTTATGGGGAGGTGTGGAAG |
| chr12_VP2 | AGAATTAACACAGAGGGGTC |
| chr12_VP3 | AAGGATTCCAGAGTCTCAGC |
| chr18_VP1 | AATAGGGGTCTCAGGGCTCT |
| chr18_VP2 | GTTCCCAACCACCTCACACC |
| chr18_VP3 | ACACTTGGCCCCGGAGGCTG |
| chr7_H19 | TTTCGGTGGACGCACGCACG |
| chr7_Igf2 | CATTGGAAGAGAGTGTGTAC |

<u>Primers</u>

***RT-qPCR primers:***

| **Target** | **Sequences (5'-3')** | |
|---|---|---|
| Site 1 (upstream of motif 1) | Fwd: | CAAACGTGCAAAATAATGTGG |
| | Rev: | GCTACCCCTTTTATCTGACC |
| Site 2 (in between motif 3 and 4) | Fwd: | GGCTGTCAGATTAACGTACT |
| | Rev: | CTGTATTTGACCCAGAAGCT |
| CTCF *Igf2* (control) | Fwd: | CACCTCAGTCGCCAAATGGT |
| | Rev: | GACTCCGCTCTGGAGGTTCT |

***4C-seq primers:***

| **Viewpoint** | **Sequences (5'-3')** |
|---|---|
| chr13_VP1 | Fwd:<br>AATGATACGGCGACCACCGAGATCTACACTCTTTCCCTACACGACGCTCTTCCGAT<br>CTAGAGAACATTAGGAAAGGCG |
| | Rev:<br>CAAGCAGAAGACGGCATACGAGATCGTGATGTGACTGGAGTTCAGACGTGTGCTCT<br>TCCGATCTACTATGGAACTTAGGCTTGAC |
| chr13_VP2 | Fwd:<br>AATGATACGGCGACCACCGAGATCTACACTCTTTCCCTACACGACGCTCTTCCGAT<br>CTGCACTGACTTGGCAAGATAT |
| | Rev:<br>CAAGCAGAAGACGGCATACGAGATCGTGATGTGACTGGAGTTCAGACGTGTGCTCT<br>TCCGATCTCACTGCTATACACACCTCTG |
| chr13_VP3 | Fwd:<br>AATGATACGGCGACCACCGAGATCTACACTCTTTCCCTACACGACGCTCTTCCGAT<br>CTACCTCATCTACTCCATAGGC |
| | Rev:<br>CAAGCAGAAGACGGCATACGAGATCGTGATGTGACTGGAGTTCAGACGTGTGCTCT<br>TCCGATCTCAGACCAGGCTATAACAAGG |

<u>ELF-Clamp: biotinylated probes for T7 promoter fusion</u>

*The first base in each probe is biotinylated*

| Viewpoint | Sequences (5'-3') |
|---|---|
| chr13_VP1 | Upstream probe:<br>TTAGACCTGCAGGTTAGATAATACGACTCACTATAGGGAGCCATTATCACAGTCCCTAAGGGAGCCCAAC |
| | Downstream probe:<br>TTAGACCTGCAGGTTAGATAATACGACTCACTATAGGGAGGTCAGCAGCAGCCTTTCAGTTTGCCAATCA |
| chr13_VP2 | Upstream probe:<br>TTAGACCTGCAGGTTAGATAATACGACTCACTATAGGGAGCTTCAGGCAGAAGTTCAACCTCGCTAGGCA |
| | Downstream probe:<br>TTAGACCTGCAGGTTAGATAATACGACTCACTATAGGGAGCGCTGGCCACACTGGGGACAGGCATAGGGC |
| chr13_VP3 | Upstream probe:<br>TTAGACCTGCAGGTTAGATAATACGACTCACTATAGGGAGCCGAGGACTGCCTTGCAGGCGTAGGAGCTG |
| | Downstream probe:<br>TTAGACCTGCAGGTTAGATAATACGACTCACTATAGGGAGCGCTGGGTGTCCATAAGCAGCCATTTTGCC |
| chr3_VP1 | Upstream probe:<br>TTAGACCTGCAGGTTAGATAATACGACTCACTATAGGGAGGTCTGCCAAGCAACTCTGCTGAAGCAGCAC |
| | Downstream probe:<br>TTAGACCTGCAGGTTAGATAATACGACTCACTATAGGGAGCTGGGGCTTCCAGAGATGGGCCCCCCAGCT |
| chr3_VP2 | Upstream probe:<br>TTAGACCTGCAGGTTAGATAATACGACTCACTATAGGGAGGAGCAGGAACGGCCCCTTTCCGTTGGGGTA |
| | Downstream probe:<br>TTAGACCTGCAGGTTAGATAATACGACTCACTATAGGGAGACAAGGCCTGCGAAAGCTCTCCTCAGGACC |
| chr3_VP3 | Upstream probe:<br>TTAGACCTGCAGGTTAGATAATACGACTCACTATAGGGAGTGTAGGCCACATCTTCACTGTAGAGGTTCA |
| | Downstream probe:<br>TTAGACCTGCAGGTTAGATAATACGACTCACTATAGGGAGGACACTTCTGCTAGCATTCTGGGCTTCAAC |
| chr3_VP4 | Upstream probe:<br>TTAGACCTGCAGGTTAGATAATACGACTCACTATAGGGAGGGCTGGACCCATAATCTATAGTGTTCACAC |
| | Downstream probe:<br>TTAGACCTGCAGGTTAGATAATACGACTCACTATAGGGAGAAAGCCTCACACGATGGGGATGTTCCAGGA |
| chr12_VP1 | Upstream probe:<br>TTAGACCTGCAGGTTAGATAATACGACTCACTATAGGGAGCCACACCTCCCCATAAGTCACTTCGCACTG |
| | Downstream probe:<br>TTAGACCTGCAGGTTAGATAATACGACTCACTATAGGGAGAAGTGGCAGGCAATAGCCCAGTGCTTTGAG |
| chr12_VP2 | Upstream probe:<br>TTAGACCTGCAGGTTAGATAATACGACTCACTATAGGGAGCCCTCTGTGTTAATTCTCCTTAAGCTT |
| | Downstream probe:<br>TTAGACCTGCAGGTTAGATAATACGACTCACTATAGGGAGGTCAGGGAGAGACCTGGATGTCCTGCTGGT |
| chr12_VP3 | Upstream probe:<br>TTAGACCTGCAGGTTAGATAATACGACTCACTATAGGGAGGAGACTCTGGAATCCTTTCCATACTTACTT |
| | Downstream probe:<br>TTAGACCTGCAGGTTAGATAATACGACTCACTATAGGGAGAGCTGGCATTTTAAAAACGGTACCCCATTT |
| chr18_VP1 | Upstream probe:<br>TTAGACCTGCAGGTTAGATAATACGACTCACTATAGGGAGTCTAGGCTGCTTGTGGGGTAGATAAGTACA |
| | Downstream probe:<br>TTAGACCTGCAGGTTAGATAATACGACTCACTATAGGGAGGCCCTGAGACCCCTATTCTTTAGTCCAGGT |
| chr18_VP2 | Upstream probe:<br>TTAGACCTGCAGGTTAGATAATACGACTCACTATAGGGAGGTGAGGTGGTTGGGAACTGCAAGCCAAACC |
| | Downstream probe:<br>TTAGACCTGCAGGTTAGATAATACGACTCACTATAGGGAGACCTGGGGACAAGGCAGTATGCGTGCTCGC |
| chr18_VP3 | Upstream probe:<br>TTAGACCTGCAGGTTAGATAATACGACTCACTATAGGGAGCTGTGGCCTCTTCTGCAATGAACCTGCTGG |
| | Downstream probe:<br>TTAGACCTGCAGGTTAGATAATACGACTCACTATAGGGAGCCTCCGGGGCCAAGTGTGTGCTCACCCACA |
| chr7_H19 | Upstream probe:<br>TTAGACCTGCAGGTTAGATAATACGACTCACTATAGGGAGGCGTGCGTCCACCGAAACCCCATAGCCATA |
| | Downstream probe:<br>TTAGACCTGCAGGTTAGATAATACGACTCACTATAGGGAGACGCGGCAGTTTCTATGTCTCCCGCCTATA |
| chr7_Igf2 | Upstream probe:<br>TTAGACCTGCAGGTTAGATAATACGACTCACTATAGGGAGCACACTCTCTTCCAATGTGATGCCCAGTGG |
| | Downstream probe:<br>TTAGACCTGCAGGTTAGATAATACGACTCACTATAGGGAGTACAGGGAGGGGCCAATGACTGCCTAGGTT |
